# Three enzymes and one substrate; regulation of flux through the glyoxylate shunt in the opportunistic pathogen, *Pseudomonas aeruginosa*

**DOI:** 10.1101/318345

**Authors:** Audrey Crousilles, Stephen K. Dolan, Paul Brear, Dimitri Y. Chirgadze, Martin Welch

## Abstract

The glyoxylate shunt bypasses the oxidative decarboxylation steps of the tricarboxylic acid (TCA) cycle, thereby conserving carbon skeletons for biosynthesis. The branchpoint between the TCA cycle and the glyoxylate shunt is therefore widely considered to be one of the most important junctions in the whole of microbial metabolism. In *Escherichia coli*, AceK-mediated phosphorylation and inactivation of the TCA cycle enzyme, *iso*citrate dehydrogenase (ICD), is necessary to redirect flux through the first enzyme of the glyoxylate shunt, *iso*citrate lyase (ICL). In contrast, Mycobacterial species lack AceK and employ a phosphorylation-insensitive *iso*citrate dehydrogenase (IDH) at the branchpoint. Flux partitioning here is controlled “rheostatically” through cross-activation of IDH by the product of ICL activity, glyoxylate. However, the opportunistic human pathogen, *Pseudomonas aeruginosa*, expresses IDH, ICD, ICL and AceK. Here, we present the structure, kinetics and regulation of each branchpoint enzyme. We show that flux partitioning is coordinated through reciprocal regulation of the enzymes involved, beautifully linking carbon flux with the availability of key gluconeogenic precursors in a way that cannot be extrapolated from an understanding of the branchpoint enzymes in other organisms.

---

The glyoxylate shunt is an anaplerotic pathway which bypasses the oxidative decarboxylation steps of the TCA cycle, thereby conserving carbon skeletons for gluconeogenesis and biomass production^1^. The mechanisms controlling carbon flux partitioning between the TCA cycle and glyoxylate shunt were largely worked out in the *Escherichia coli* model in the 1980s. Here, the TCA cycle enzyme, *iso*citrate dehydrogenase (ICD, encoded by *icd*), and the glyoxylate shunt enzyme, *iso*citrate lyase (ICL, encoded by *aceA*) compete for available *iso*citrate. ICD has a much lower K_M_ for *iso*citrate (K_M_ 8 μM^2^) than ICL (K_M_ 604 μM^3^), so in order to get significant flux through the glyoxylate shunt, ICD needs to be inactivated. This is accomplished by reversible phosphorylation of ICD on Ser 113, mediated by a dual function kinase/phosphatase, AceK^4^. Phosphoserine 113 is thought to electrostatically repulse *iso*citrate, thereby preventing substrate binding^5^. Consequently, when the kinase activity of AceK is dominant, ICD becomes phosphorylated and inactivated, allowing carbon flux to be redirected through the glyoxylate shunt. In contrast, when the phosphatase activity of AceK dominates, flux is restored through the TCA cycle. The ratio of kinase:phosphatase activity in AceK is controlled by allosteric regulators^6^.

Not all bacteria share the same enzymology as *E. coli* at the TCA cycle-glyoxylate shunt branchpoint (“TGB”); some bacteria encode a second, AceK-insensitive *iso*citrate dehydrogenase isozyme, IDH. Genome sequencing efforts have revealed that some species, such as the industrially-important fermenter *Corynebacterium glutamicum*, contain only *idh*, whereas others contain only *icd* (e.g., *E. coli*). Generally, and as might be expected, there is a strong correlation between the presence of genes encoding ICD, ICL and AceK in a genome. There are exceptions to this though. For example, *Mycobacterium tuberculosis* encodes ICL, ICD and IDH, but lacks AceK. Without a mechanism to inactivate ICD, this raises the question of how flux is redirected through the glyoxylate shunt in this organism. This was partially resolved by the finding that ICD appears to be fully dispensable for flux through the TCA cycle in *M. tuberculosis*^7^, so only ICL and IDH compete for the pool of *iso*citrate. Recent work done in a related species of mycobacteria (*M. smegmatis*) indicates that flux partitioning at the TGB is determined primarily by allosteric activation of IDH by ICL-derived glyoxylate^7^.

An additional layer of unexplored complexity is seen in the opportunistic human pathogen, *Pseudomonas aeruginosa.* Here, <u>three</u> enzymes compete for *iso*citrate; ICD, IDH and ICL. Moreover, *P. aeruginosa* also encodes the ICD kinase/phosphatase, AceK. This raises the question of how AceK-mediated post-translational regulation of ICD activity is integrated with the activities of ICL and IDH. The lack of clarity here is ironic, especially given that Kornberg and Krebs originally elucidated the glyoxylate shunt in a *Pseudomonas* strain, KB1^8^. Therefore, and to investigate the enzymology of the TGB further, we cloned and purified ICL, ICD, IDH and AceK from *P. aeruginosa*. Kinetic analysis of the purified enzymes revealed that in striking contrast with *E. coli* and *M. smegmatis*, *P. aeruginosa* ICL has a *higher* affinity for *iso*citrate (K_M_ 12 μM) compared with the competing dehydrogenases, ICD (K_M_ 26 μM) and IDH (K_M_ 18 μM); **Table I** and **Figure S1**. ICD and IDH shared similar K_M_ values for their other substrate, NADPH (32 μM and 34 μM, respectively). ICD and IDH were expressed at comparable levels during growth on either glucose or acetate (**Figure S1E**). Commensurate with earlier observations that there is significant flux through the glyoxylate shunt even during growth on glucose^9^, ICL was expressed (albeit at low levels) on this substrate. However, ICL expression increased during growth on acetate. Again, this is consistent with earlier observations that acetate induces expression of the ICL-encoding gene, *aceA*^10^.

**Table I:** Kinetic parameters of ICL, IDH and ICD.

| | $K_M$<br>($\mu M$) | $V_{max}$<br>($\Delta mM/s$) | $k_{cat}$ ($s^{-1}$)<sup>1</sup> | $k_{cat}/K_M$<br>($s^{-1}/\mu M$) | $K_i$ (mM) | Comments |
| --- | --- | --- | --- | --- | --- | --- |
| **ICL** |  |  |  |  |  |  |
| *isocitrate* | 12.2 | 0.8 | 18.8 | 1.6 | n/a | n/a |
| *citrate* | 18.8 | 0.8 | 17.7 | 0.9 | 15.8 | Non-competitive |
| *coenzyme A* | 22.6 | 0.6 | 14.9 | 0.7 | 3.9 | Non-competitive |
| *glycolic acid* | 18.2 | 0.5 | 12.2 | 0.7 | 1.8 | Non-competitive |
| *itaconic acid* | 6.3 | 0.4 | 8.7 | 1.4 | 0.9 | Uncompetitive |
| *maleic acid* | 3.6 | 0.3 | 6.3 | 1.8 | n/a | Mixed |
| *malic acid* | 16.6 | 0.7 | 15.3 | 0.9 | 4.4 | Non-competitive |
| *malonic acid* | 12.0 | 0.5 | 12.2 | 1.0 | 2.9 | Uncompetitive |
| *nitropropionic acid* | 6.9 | 0.3 | 7.5 | 1.1 | 1.0 | Uncompetitive |
| *oxaloacetate* | 18.4 | 0.5 | 11.0 | 0.6 | 1.4 | Non-competitive |
| *phosphoenolpyruvate* | 6.0 | 0.4 | 8.9 | 1.5 | n/a | Mixed |
| *pyruvate* | 18.2 | 0.5 | 11.8 | 0.7 | 1.7 | Non-competitive |
| *succinate* | 7.8 | 0.3 | 7.1 | 0.9 | 0.9 | Uncompetitive |
| **IDH** |  |  |  |  |  |  |
| *isocitrate* | 17.7 | 0.5 | 183.7 | 10.4 | n/a | n/a |
| *glyoxylate* | 20.7<sup>2</sup> | 2.3 | 925.2 | 44.8 | n/a | Non-competitive |
| *oxaloacetate* | 39.2<sup>2</sup> | 3.0 | 1210.9 | 30.9 | n/a | Uncompetitive |
| *pyruvate* | 42.0<sup>2</sup> | 3.5 | 1408.2 | 33.6 | n/a | Uncompetitive |
| NADP | 34.5<sup>2</sup> | 0.5 | 210.9 | 6.1 | n/a | n/a |
| *glyoxylate* | 160.4<sup>2</sup> | 3.8 | 1544.2 | 9.6 | n/a | Uncompetitive |
| *oxaloacetate* | 29.0<sup>2</sup> | 3.0 | 1217.7 | 42.0 | n/a | Non-competitive |
| *pyruvate* | 49.5 | 3.4 | 1381.0 | 27.9 | n/a | Non-competitive |
| **ICD** |  |  |  |  |  |  |
| *isocitrate* | 25.6 | 1.3 | 608.8 | 23.8 | n/a | n/a |
| α-ketoglutarate | 64.0 | 1.6 | 745.8 | 11.7 | 666.0 | Competitive |
| NADP | 32.3 | 1.5 | 684.9 | 21.2 | n/a | n/a |

$K_M$ and $V_{max}$ values were calculated after fitting to the Michaelis-Menten curve in Prism 6 GraphPad. Values of $k_{cat}$ were derived from $V_{max}$ . <sup>1</sup>The value of $E_t$ was based on the most likely oligomeric status of the native protein from the combined crystallographic and solution structure data. ICL was assumed to be a tetramer, ICD a dimer, and IDH a monomer. <sup>2</sup>Regulators of IDH altered the kinetics from hyperbolic to sigmoidal with respect to *isocitrate*, so the reported values in the Table are $K_{half}$ , where $h$ is the Hill coefficient, instead of $K_M$ . The value of $K_i$ was determined differently depending on the type of inhibition observed. In the case of competitive inhibition, we used $K_i = \frac{K_M \times [I]}{K_{mobs} - K_M}$ , whereas for non-competitive inhibition, we used $K_i = \frac{[I]}{(V_m/V_{mobs}) - 1}$ . For uncompetitive inhibition, $K_i'$ was determined from the $y$ -axis intercept of the Lineweaver-Burk plot, assuming that $K_i' = \frac{[I]}{(y_{intercept} \times V_{max}) - 1}$ . In all the cases, $[I] = 1 mM$ .

## Structure and regulation of *P. aeruginosa* ICL

At 531 amino acids, *P. aeruginosa* ICL is longer than *E. coli* ICL (434 amino acids) and shares only 27% amino acid sequence identity with the latter. It also lies on a distinct branch of the evolutionary tree for this enzyme (which also includes ICL from pathogens such as *Burkholderia cepacia* and *Acinetobacter baumanii*), with no structurally-characterised homologues to date (**Figure S2**). To investigate this further, we solved the x-ray crystal structure of *P. aeruginosa* ICL to 1.9Å resolution, with Ca^2+^ and glyoxylate embedded in the active site. Crystallographic statistics are given in **Table S1**. The presence of a Ca^2+^ (which was coordinated with the glyoxylate ligand and three water molecules) was confirmed using Checkmymetal. In the crystal structure, there is one ICL polypeptide per asymmetric portion of the unit cell, which belonged to the I222 space group (**Figure 1A**). These polypeptides are related by a crystallographic two-fold axis to yield a tetrameric structure (**Figure 1B**) with extensive contacts made between each protomer (**Figure S3**). This is consistent with gel filtration/multi-angle light scattering (GFC/MALS) and analytical ultracentrifugation (AUC) data indicating that ICL behaves as a 231 kDa tetramer in solution (**Figure S4**). Each protomer in *P. aeruginosa* ICL is comprised of 17 α-helices and 14 β-strands arranged around a central TIM barrel-like core (secondary structure assignments are shown in **Figure S5**). However, the overall fold of each protomer is distinct when compared with ICL from *E. coli* and *Mycobacterium tuberculosis* (**Figure 1C**). In particular, helices α1 and α2 of *P. aeruginosa* ICL are flipped as a unit by almost 180° relative to the axis of α3, and α6 is extended by an additional 4 helical turns. In addition, α13 and α14 extend away from the globular core of the protein, generating structural projections which give the tetramer a distinctly rugged, star-like profile. The most obvious difference though, is the presence of a relatively unstructured “head-domain” spanning the region between Ile272 and Ile306. The head domains of adjacent protomers form some of the more intimate contacts underpinning tetramer formation (**Figure 1B** and **Figure S3**). Interestingly, ICL from the fungus *Aspergillus nidulans* also has a head-like domain, although this is rich in α-helices and does not appear to play a significant role in inter-protomer interactions^11^.

**Figure 1.**
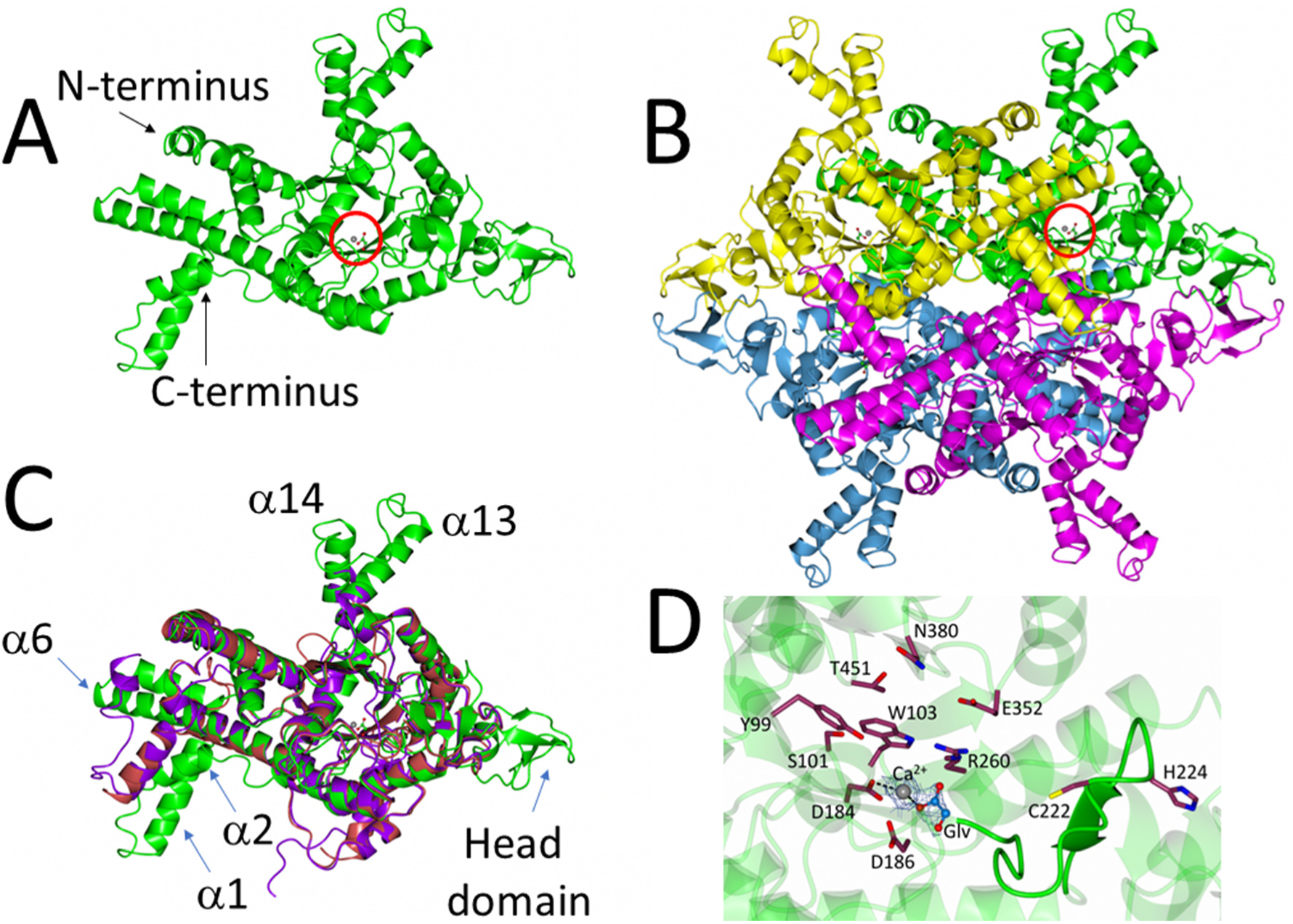
Structure of *P. aeruginosa* ICL. **(A)** Cartoon representation of a single *iso*citrate lyase protomer from *P. aeruginosa* (PDB 6G1O). The bound Ca^2+^ and glyoxylate are indicated with a red circle. **(B)** Quaternary structure of the *iso*citrate lyase in the crystal. Note the β strand-rich head domains on either side of the tetramer and the α helical projections (α13 and α14 in each protomer) above and below the structure. The Ca^2+^ and glyoxylate in one of the protomers are circled. **(C)** Superposition of *P. aeruginosa* ICL (green), *M. tuberculosis* ICL (purple, PDB 1F8I) and *E. coli* ICL (red, PDB 1IGW) revealing that the catalytic core of the enzyme is highly conserved. Consistent with this, the root mean square deviation (RMSD) between *P. aeruginosa* and *E. coli* ICL is 1.35 Å, and the RMSD between *P. aeruginosa* and *M. tuberculosis* ICL is 1.30 Å. However, note how α1 and α2 are flipped by 180° in *P. aeruginosa* ICL, the extended helices α13 and α14, and the presence of a head-domain (absent from *M. tuberculosis* and *E. coli* ICL. **(D)** Conserved active site residues in *P. aeruginosa* ICL. The β4-β5 loop containing the general base, Cys222, is highlighted. The electron density map for glyoxylate and Ca^2+^ is contoured at 1.7σ.

The catalytic core of the enzyme is conserved. The active site of ICL is comprised of two distinct parts; those residues associated with the β-strands and loops around the rim of the TIM barrel, and a flexible loop of structure between β4 and β5^12^. The latter contains the catalytic general base, Cys222 (*P. aeruginosa* numbering). In our structure, this β4-β5 loop points away from the active site. Presumably, in the presence of *iso*citrate, this loop swings inwards to cap the active site, as it does in ICL from *M. tuberculosis*^13^. The glyoxylate moiety in our structure is displaced by 6Å towards the β4-β5 loop, leaving it poised ready to exit the active site, although it remains anchored on the enzyme through coordination with the Ca^2+^ (**Figure 1D**).

Given its position at the TGB, it is unsurprising that the activity of *P. aeruginosa* ICL is modulated by certain metabolites. *P. aeruginosa* ICL activity was inhibited by oxaloacetate, pyruvate, succinate, phospho*enol*pyruvate and CoA (**Figure 2**). No activators were identified. Oxaloacetate (K_i_ 1.9 mM), pyruvate (K_i_ 2 mM) and CoA (K_i_ 1.2 mM) all inhibited the enzyme non-competitively (**Figure 2A**) whereas the reaction product, succinate, inhibited uncompetitively (K_i_’ 0.6 mM) and phospho*enol*pyruvate (PEP) displayed mixed inhibition (**Figure 2B**). In common with other *iso*citrate lyases, the enzyme was also susceptible to uncompetitive inhibition by itaconic acid (K_i_’ 0.5 mM) and nitropropionic acid (K_i_’ 0.6 mM)^14,15^, and potent mixed inhibition by maleic acid (**Figure 2C**). The latter may reflect the partial structural similarity between PEP and maleic acid.

**Figure 2.**
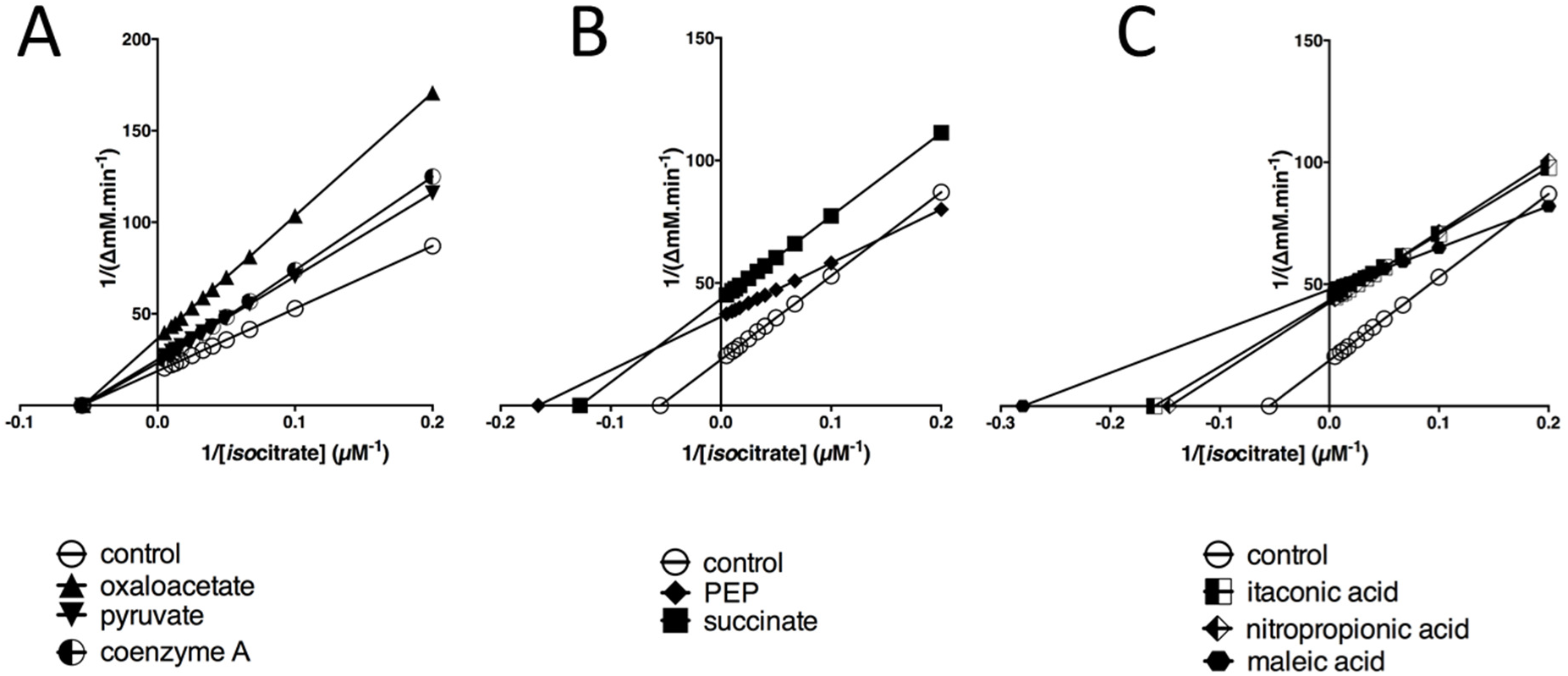
Inhibition of *P. aeruginosa* ICL activity by small molecules. **(A)** Coenzyme A, oxaloacetate and pyruvate non-competitively inhibit ICL. The figure shows a Lineweaver-Burk transformation of ICL kinetic data obtained in the presence of no addition (control) or in the presence of oxaloacetate, pyruvate, or coenzyme A. **(B)** Lineweaver-Burk plot showing that succinate inhibits ICL uncompetitively whereas phospho*enol*pyruvate displays mixed inhibition. **(C)** Itaconate and nitropropionate inhibit ICL uncompetitively, whereas maleic acid however displays mixed inhibition. The plots were generated using GraphPad Prism 6. All small molecules (except the substrate) were added at 1 mM final concentration.

## Structure and regulation of *P. aeruginosa* IDH

In *P. aeruginosa*, two *iso*citrate dehydrogenases, IDH and ICD, compete with ICL for *iso*citrate. In solution, IDH (monomeric molecular mass 82 kDa) had an apparent molecular mass of 235 kDa (GFC/MALS) or 273 kDa (AUC), suggesting that it adopts a higher-order structure (**Figure S6**). To investigate the enzyme further, we solved its x-ray crystal structure to 2.7Å resolution (**Table S1**). The asymmetric unit comprised two molecules of IDH; one (designated chain A) contained bound α-*keto*glutarate and NADP, and the other (chain B) contained no bound substrate/product (**Figure 3A**). However, the interface between these monomers was small and analysis using PISA^16^ suggested that the two molecules cannot be considered to be protomers of a functional dimer. Comparison of the three-dimensional structure of the *P. aeruginosa* A and B chains revealed large conformational differences, especially in the smaller domain of each chain (**Figure S7**), suggesting that catalysis is accompanied by structural rearrangements.

**Figure 3.**
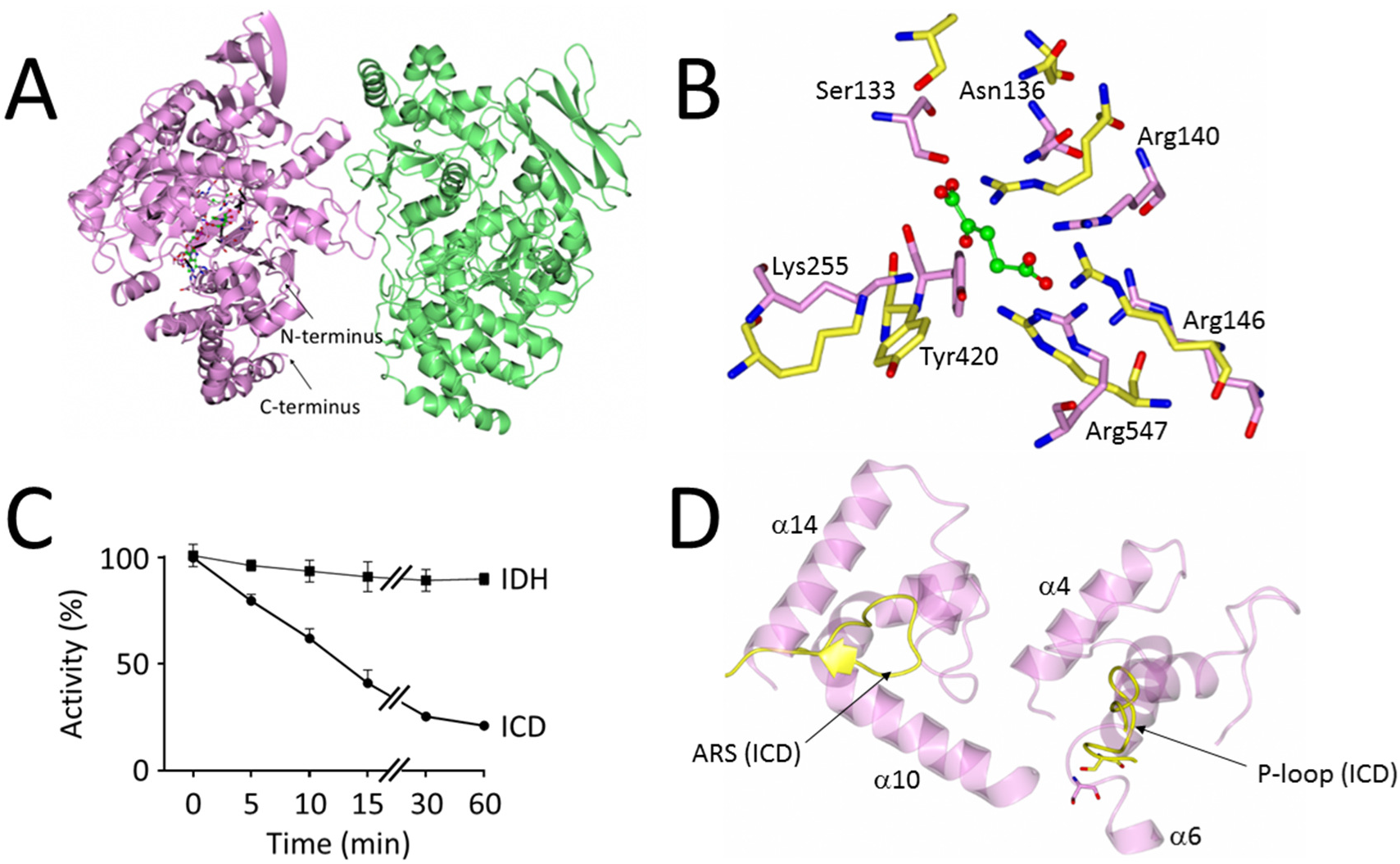
Structure and regulation of *P. aeruginosa* IDH. **(A)** Cartoon schematic of *iso*citrate dehydrogenase (IDH) from *P. aeruginosa*. Chain A (mauve) contains one molecule each of NADP^+^ and α-*keto*glutarate bound in its active site. Chain B (light green) does not contain any bound small molecules. The two IDH chains in the asymmetric unit do not show enough inter-protein contacts to warrant being labelled as protomers in a dimer. **(B)** Close-up view of the conserved active site residues in *P. aeruginosa* IDH (light pink, PDB 6G3U) and *P. aeruginosa* ICD (yellow, PDB 5M2E). Residue numbering is based on the IDH sequence. **(C)** *P. aeruginosa* ICD, but not IDH, is inactivated by AceK-dependent phosphorylation. The figure shows the loss of *iso*citrate dehydrogenase activity over time following treatment with AceK/ATP. Reaction mixtures (200 μL) contained 100 mM Tris-HCl (pH 7.0), 1 mM ATP, 2 mM MgCl_2_, 5 μg purified *P. aeruginosa* AceK, and 10 μg *P. aeruginosa* ICD or IDH (as indicated). Reactions were allowed to proceed at 37°C, and at the indicated times, aliquots were withdrawn and assayed immediately for *iso*citrate dehydrogenase activity as described in Materials and Methods. Activity was considered to be 100% at T0. (D) The architecture of the active site is different in ICD and IDH. In spite of the conserved constellation of active site residues (panel (B)) in *P. aeruginosa* ICD and IDH, the P-loop containing the phosphoserine in ICD (yellow) and the AceK recognition sequence (ARS) are replaced in IDH by two helices α10-α14 and α4-α6, respectively. This altered arrangement prevents AceK from accessing the active site serine in IDH.

The α-*keto*glutarate binding cleft is sandwiched between the two IDH domains (**Figure S7A**), with side chains from both domains involved in binding. The active site residues are remarkably well-conserved between IDH and ICD (**Figure 3B**). This is consistent with the proposed common evolutionary origins of both proteins^17^, and raises the question of why IDH is not a substrate for AceK (**Figure 3C**). Comparison of the *P. aeruginosa* IDH active site architecture with that of ICD in the AceK-ICD complex from *E. coli* (PDB; 3LCB) reveals that in spite of the conserved constellation of catalytic residues, the active sites are differently-structured, with the two ICD motifs known to be critical for recognition by AceK (designated the P-loop and <u>A</u>ceK <u>r</u>ecognition <u>s</u>egment (ARS)) absent in IDH (**Figure 3D**).

Glyoxylate was recently reported to be the principal regulator of IDH in *M. smegmatis*, enabling “rheostatic” control of flux through the glyoxylate shunt^7^. However, and although glyoxylate did stimulate *P. aeruginosa* IDH activity, pyruvate and oxaloacetate were far more potent regulators. All three compounds changed the *iso*citrate dependency of IDH kinetics from hyperbolic to sigmoidal (**Figure 4A**), and all had a pronounced effect on k_cat_ with only a small impact on K_M_ (**Table I**), indicating that the enzyme is of the rarer V-type allosteric class. The shift to sigmoidal *iso*citrate kinetics in the presence of these activators suggests cooperativity in substrate binding, consistent with IDH adopting a higher-order structure in solution. However, these effectors had much less impact on the NADP-dependency of IDH kinetics, which remained essentially hyperbolic (**Figure 4B**). In common with previously characterised *iso*citrate dehydrogenases, IDH (and also, ICD) was strongly inhibited by mixtures containing oxaloacetate and glyoxylate, presumably due to the non-enzymatic formation of oxalomalate^18^.

**Figure 4.**
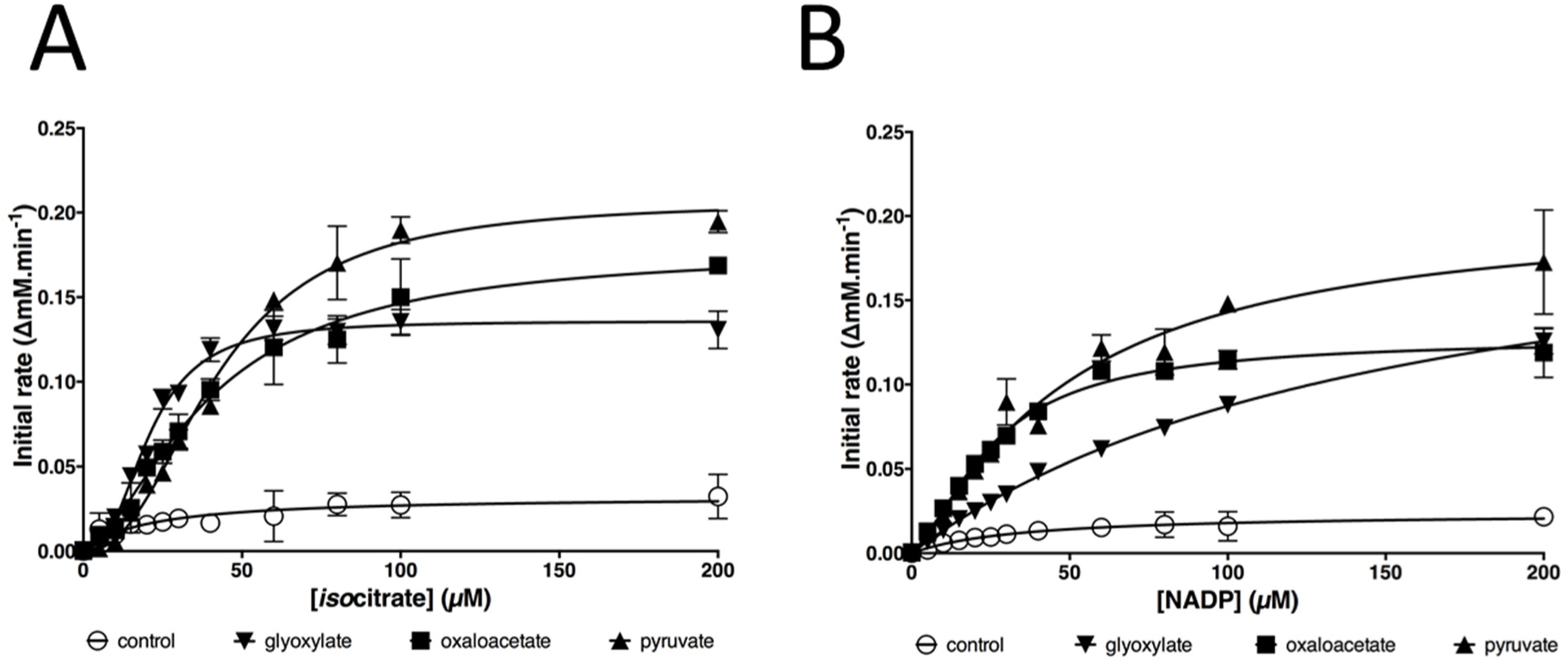
Pyruvate, oxaloacetate and glyoxylate strongly activate the *iso*citrate dehydrogenase activity of IDH. **(A)** Effect of pyruvate, oxaloacetate and glyoxylate of the *iso*citrate-dependency of IDH kinetics. **(B)** Effect of pyruvate, oxaloacetate and glyoxylate of the NADP^+^-dependency of IDH kinetics. Each regulator was added at 1 mM final concentration. Error bars correspond to ± 1 standard deviation from n = 3 replicates.

## Structure and regulation of ICD

The second *iso*citrate dehydrogenase encoded by *P. aeruginosa* is ICD. *P. aeruginosa* ICD crystallised as a dimer (**Figure S8, Table S1**) and also behaved as a dimer in solution (**Figure S9**). The structure, solved to 2.7Å resolution, was very similar to that reported for the *E. coli* enzyme (**Figure S8B**), with a clasp-like structure mediating the dimerization^19^. The residues lining the substrate binding site were also conserved between the *E. coli* and *P. aeruginosa* enzymes (**Figure S8C**). ICD was competitively inhibited by the reaction product, α-*keto*glutarate (K_i_ 666 μM, **Figure S8D**). However, the main mode of regulation was through AceK-dependent phosphorylation (**Figure 5A,B**). Interestingly, in all of our experiments with AceK and ICD from *P. aeruginosa* (hereafter, AceK_PA_ and ICD_PA_), the maximal inhibition attributable to AceK-mediated phosphorylation was only around 75% (**Figure 5B**). This is not due to a lower intrinsic activity of AceK_PA_, because AceK_PA_ was able to almost completely inactivate ICD from *E. coli* (ICD_EC_) (**Figure 5B**). Comparison of the ICD_PA_ and ICD_EC_ active site region revealed slight differences in the spatial disposition of the P-loop required for AceK recognition (**Figure S8E**), making it formally possible that ICDPA is less sensitive to AceK-mediated phosphorylation. However, AceK_EC_ was able to completely inactivate ICD_PA_, indicating that these local differences in P-loop conformation are unlikely to be functionally significant (**Figure 5B**). Taken together, these data suggest that in the absence of external factors to stimulate its kinase activity, AceK_PA_ cannot fully inhibit ICD_PA_.

**Figure 5.**
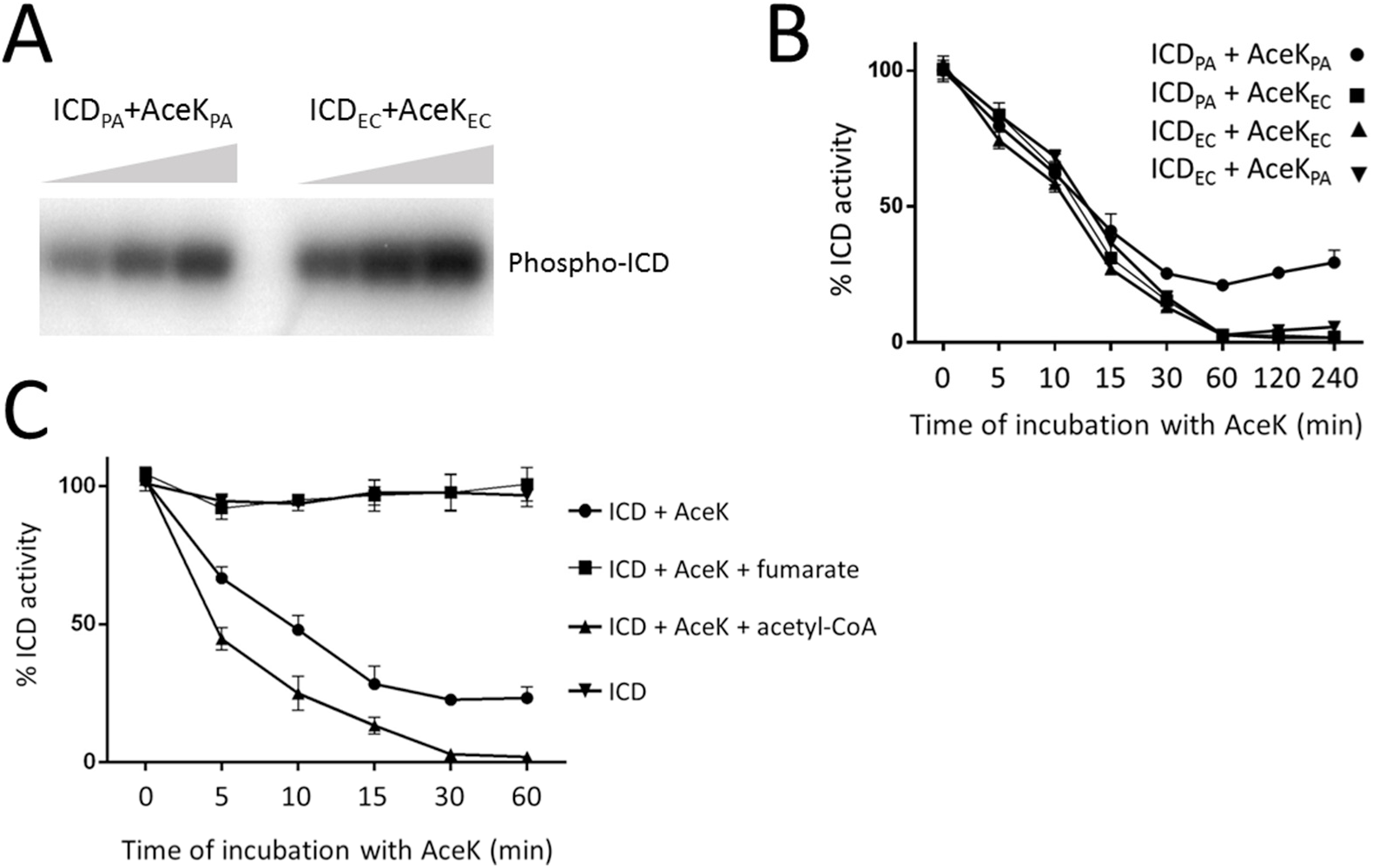
Regulation of *P. aeruginosa* ICD activity by AceK-dependent phosphorylation. **(A)** AceK-mediated phosphotransfer from γ[^32^P]ATP to ICD. Phosphotransfer was measured between *P. aeruginosa* AceK and *P. aeruginosa* ICD (AceK_PA_ and ICD_PA_) or between *E. coli* AceK and *E. coli* ICD (AceK_EC_ and ICD_EC_), as indicated. Aliquots were removed for sampling after 1, 2 and 3 min incubation, and resolved by SDS-PAGE. Radioactivity was monitored using a Phosphoimager. **(B)** *P. aeruginosa* AceK is intrinsically less efficient at phosphorylating *P. aeruginosa* ICD than *E. coli* AceK. The figure shows a “mix n’ match” experiment in which the efficiencies of *P. aeruginosa* and *E. coli* AceK at inactivating the ICD homologues from each species are measured. ICD_PA_ and ICD_EC_ can each be phosphorylated by either AceK_PA_ or AceK_EC_. However, the ICD_PA_/AceK_PA_ pair never achieves full inactivation, in spite of the fact that AceK_PA_ fully inactivates ICD_EC_ (indicating that the kinase is competent to do this) and that AceK_EC_ fully inactivates ICD_PA_ (indicating that the ICD is potentially fully inactivatable). Reaction conditions as in Figure 3. **(C)** Fumarate stimulates the phosphatase activity of AceK_PA_ (thereby reactivating ICD), whereas acetyl-CoA stimulates the kinase activity of AceK_PA_ (thereby inactivating ICD). Fumarate and acetyl-CoA were present at 5 mM concentration.

Although the *P. aeruginosa* and *E. coli* ICD enzymes are 79% identical at the amino acid sequence level, and the site of phosphorylation (Ser115 in *P. aeruginosa* ICD) is conserved, the corresponding AceK orthologues show substantial differences. In particular, the N-terminal regulatory domain is just 35% identical between the two species. In contrast, the C-terminal catalytic domain is 60% identical and retains all the sequence motifs thought to be important for kinase and phosphatase activity. These data suggest that *P. aeruginosa* AceK is likely to be regulated differently compared with the *E. coli* orthologue. To investigate this further, we examined how a panel of potential regulators affected AceK_PA_-dependent inactivation of ICD_PA_ (**Figure S10**). ICD was first phosphorylated by AceK for 60 min (enough to maximally inactivate the dehydrogenase (**Figure 5B**)). Following this, the indicated potential regulators were added and restoration of *iso*citrate dehydrogenase activity was monitored after 30 min (**Figure S10A**). As a control, and to confirm the species specificity of each regulator, we also examined (*i*) whether the same compounds affected AceK_EC_-dependent restoration of ICD_EC_ activity (**Figure S10B**) and (*ii*) whether the regulators had any intrinsic impact on ICD activity (**Figure S11**). With the exception of citrate (which presumably competitively blocks the ICD active site) and to a lesser extent, glyceraldehyde-3-phosphate and phospho*enol*pyruvate, none of the tested potential regulators affected ICD activity directly. The phosphatase activity of AceK_EC_ was stimulated by α-*keto*glutarate, glyceraldehyde 3-phosphate, 2-*keto*-3-deoxyphosphogluconate (KDPG), pyruvate and oxaloacetate, and to a much lesser extent by fumarate, AMP and ADP (**Figure S10B**). The phosphatase activity of AceK_PA_ was stimulated by an overlapping, but larger number of regulators, including α-*keto*glutarate, fructose 1,6-bisphosphate, glyceraldehyde 3-phosphate, glyoxylate, AMP, ADP, glycolate, pyruvate and oxaloacetate. Intriguingly, fumarate was the most potent activator of the phosphatase in AceK_PA_, yet this compound only weakly stimulated the phosphatase activity of AceK_EC_. Similarly, succinate had no apparent effect on AceK_EC_ but strongly stimulated AceK_PA_ phosphatase activity. In contrast, the Entner-Doudoroff pathway intermediate, KDPG, only marginally stimulated AceK_PA_ phosphatase activity, yet this compound was one of the more potent activators of the AceK_EC_ phosphatase. Perhaps the most noticeable regulator was acetyl-CoA. This compound weakly stimulated the phosphatase activity of AceK_EC_, whereas in in AceK_PA_, it appeared to stimulate the kinase activity. To investigate this further, we conducted a time-course analysis of ICD_PA_ activity following AceK_PA_ treatment in the presence of acetyl-CoA, and fumarate (**Figure 5C**). What is immediately apparent is that, whereas fumarate maximally stimulates AceK_PA_ phosphatase activity, acetyl-CoA maximally activates the AceK kinase activity, allowing the enzyme to inactivate >75% of the ICD, and at a faster rate than AceK alone.

## Discussion

The driver behind this work is that *P. aeruginosa* exhibits a particular predilection for catabolising fatty acids, especially during infection scenarios^20^. Under these conditions, the glyoxylate shunt becomes centrally-important for growth. Indeed, a mutant of *P. aeruginosa* in which ICL and malate synthase (MS^21^), are absent is cleared from a mouse pulmonary infection model within 48 hours^22^, indicating that the shunt is an excellent target for the development of adjuvant interventions. The physiological importance of the shunt in infection may not solely be due to the metabolic defect either; *aceA* mutants also show pronounced defects in virulence^23–25^. Consequently, therapeutic agents which inhibit flux through the glyoxylate shunt (or redirect flux away from the shunt) could potentially deliver a powerful “double whammy” by eliciting both metabolic insufficiency and reduced virulence. However, a better understanding of the enzymology of the *P. aeruginosa* TGB will be essential if we are to realise its full potential as a therapeutic target. Indeed, in the current work, we show that this understanding cannot be gleaned by extrapolation from previously-characterised species. For example, in *P. aeruginosa*, three enzymes compete for *iso*citrate, and ICL has a <u>higher</u> affinity for the substrate than the competing dehydrogenases. Also, the enzymology appears to be more complex in *P. aeruginosa* compared with other species, involving both AceK (absent in e.g., *M. smegmatis* and *M. tuberculosis*) and IDH (absent in many enterics, such as *E. coli*).

The key regulators in the *P. aeruginosa* system are oxaloacetate and pyruvate, which reciprocally regulate ICL and IDH, thereby elegantly coordinating metabolic flux partitioning between the TCA cycle and glyoxylate shunt. When these compounds are abundant (signalling to the cell that there are sufficient gluconeogenic precursors for biomass production), IDH becomes activated and ICL inhibited, leading to greater flux around the TCA cycle. In a similar vein, oxaloacetate and pyruvate also stimulate the phosphatase activity of AceK, leading to increased ICD activity. In contrast, when these gluconeogenic precursors are in short supply, IDH becomes deactivated and ICL becomes disinhibited, thereby restoring flux through the glyoxylate shunt. It is worth noting that ICD activity also becomes depressed *via* a different mechanism when demand for gluconeogenesis is high. Acetyl-CoA strongly stimulates the kinase activity of AceK, leading to inactivation of ICD (again, promoting flux through the glyoxylate shunt). This is significant because acetyl-CoA would be expected to accumulate during growth on fatty acids or acetate, or when its condensation partner in the TCA cycle, oxaloacetate, is in short supply (signalling that anaplerosis is necessary). As might be expected, uncharged CoA was an inhibitor of ICL, suggesting that flux through the bifurcation point is also partially dictated by enzymatic surveillance of the acetyl-CoA : CoA ratio.

In addition to gluconeogenic precursors, we also found that the products of ICL activity *per se* play an important role in coordinating flux partitioning; succinate feeds back to inhibit ICL and also stimulates the phosphatase activity of AceK (thereby re-activating ICD). Similarly, glyoxylate also stimulates the AceK phosphatase (reactivating ICD), and directly activates IDH (although not as strongly as pyruvate or oxaloacetate). In *M. smegmatis*, this type of mechanism has been termed “rheostatic”^7^, although in *P. aeruginosa*, this appears to be secondary compared with the more dominant role(s) played by oxaloacetate, pyruvate and acetyl-CoA.

Unlike most other organisms characterised to date, the ICL in *P. aeruginosa* has a lower K_M_ for *iso*citrate than the *iso*citrate dehydrogenases. This may be to always ensure adequate carbon flux through TCA cycle, especially when demand for NADPH is high e.g., during oxidative stress or anabolism. The *iso*citrate dehydrogenase reaction is one of the main sources of NADPH in the cell^26^, and it may be that *P. aeruginosa* has evolved a “belt-and-braces” solution to ensure that it never runs out of this important anti-oxidant, even when the demand for flux through the glyoxylate shunt is high. This may also explain why it encodes two independently (albeit, similarly) regulated *iso*citrate dehydrogenases, ICD and IDH. The same logic may also explain why, even after prolonged exposure to AceK, ICD retains some activity (although this was not the case in *E. coli*, where 100% inhibition was achieved after treatment with either AceK_PA_ or AceK_EC_). Complete ICD_PA_ inhibition was only observed in the presence of acetyl-CoA, presumably because an accumulation of this metabolite indicates to the cell that the “relief valve” of the glyoxylate shunt needs to be fully opened.

Although ICD and IDH were expressed at similar levels during growth on glucose and acetate, ICL showed strong induction on the latter. Presumably, this reflects induction of the *aceA* gene, and this has been noted by earlier workers^10^. However, and unlike the situation in *E. coli*, *aceA* is expressed at appreciable levels on glucose, and there is significant flux through the glyoxylate shunt on this carbon source^27^. This may reflect the fact that (for reasons that are not yet clear), under some conditions, optimal virulence is dependent upon flux through the glyoxylate shunt^23^.

One of the more remarkable observations made in this work relates to the structure of ICL. This large protein is an established drug target^28^, and clearly responds to multiple regulators. We found that much of the significant additional sequence present in this sub-class of ICL enzymes manifests itself as structural features projecting away from the globular core of the enzyme. Although we do not yet know what these projections do, one possibility (given their surface exposure) is that they are involved in protein-protein interactions, and we are currently investigating this. Another area of ongoing interest is to understand at an atomic level of resolution how the regulator molecules exert their effects, especially on ICL and IDH. V-type allosteric regulators remain relatively poorly characterised, and a better understanding of these may open the way towards the design of drug-like “anti-activators” as well as inhibitors.

In conclusion, we have shown that the TGB in *P. aeruginosa* has a complex enzymology that is profoundly different to that in all other organisms for which the TGB has been characterised to date. This notwithstanding, common themes are discernable among the identified regulatory molecules, and the structural data lay a solid groundwork for downstream drug design efforts.

## Acknowledgements

This work was supported by BBSRC grant BB/M019411/1.

## Methods

### Cloning, Overexpression, and Purification

The ORFs encoding the ICL (PA2634), IDH (PA2624) and ICD (PA2623) enzymes were PCR-amplified from the genomic DNA of *P. aeruginosa* strain PAO1 and cloned into a derivative of the NEB vector pMAL-c2X which had been previously modified to introduce a hexa-histidine tag onto the N-terminus of the MBP. The constructs were confirmed by DNA sequencing and introduced into *E. coli* DH5α. For over-expression, the cultures were grown with good aeration in LB medium at 37°C to OD_600_ = 0.6. IPTG was then added to a final concentration of 0.3 mM and growth was allowed to continue for a further 2 h. The cells were harvested by sedimentation (3430 × *g*, 20 min, 4°C) and the pellets were resuspended in 60 mL of buffer A (200 mM NaCl, 20 mM Tris-HCl, 1 mM EDTA, 1 mM DTT, pH 7.4) containing a Complete EDTA-free protease inhibitor cocktail tablet (Roche). The cells were lysed by sonication on ice and the cell debris was removed by sedimentation (15000 × *g*, 30 min, 4°C). The supernatant was applied to an amylose resin column and washed (overnight) with 500 mL of buffer A at 4°C. The protein was then eluded in 10 mL of buffer B (200 mM NaCl, 20 mM Tris-HCl, 1 mM EDTA, 10 mM maltose, pH 7.4). The eluted sample was dialysed against 2 × 1L of buffer C (50 mM NaCl, 25 mM Tris-HCl, 10% (v/v) glycerol, pH 7.4). Following this, the sample was concentrated and cleaved with Factor Xa protease (1:100 ratio of protease:sample) at 6°C for 24h with constant gentle mixing. The protease was removed using *p*-aminobenzamidine agarose and the cleaved protein mixture was applied to a column (2 mL packed resin volume) of Ni-NTA resin equilibrated with buffer D (100 mM NaCl, 50 mM Tris-HCl, 5% (v/v) glycerol, 5 mM β-mercaptoethanol, pH 7.5) to remove the His_6_-MBP tag. The flow through (consisting of cleaved purified target protein) was collected and dialysed against buffer E (100 mM NaCl, 25 mM Tris-HCl, 10% (v/v) glycerol, 1 mM EDTA, 1 mM DTT, pH 7.5). The dialysed sample was concentrated by ultrafiltration and then snap-frozen in liquid nitrogen for storage at -80°C. Protein concentration was determined spectrophotometrically using the calculated molar extinction (ε_calc_ = 54320 M^-1^.cm^-1^ (ICL), 82280 M^-1^.cm^-1^ (IDH) and 57870 M^-1^.cm^-1^ (ICD)).

The AceK (PA1376), ICD_PA_ and ICD_EC_ (from *E. coli* strain DH5α) used for phosphotransfer/AceK-inhibition assays were purified slightly differently. The PCR-amplified ORFs were cloned into the expression vector pET-19m, which introduces a TEV-cleavable N-terminal hexa-histidine tag onto each protein. For purification of the His_6_-tagged proteins, the cells were grown in LB medium at 37°C with good aeration to OD_600_ = 0.5. The temperature was then lowered to 20°C and IPTG was added to 0.5 mM final concentration to induce expression of the cloned genes. The induced cultures were grown for a further 16 h and then harvested by sedimentation (6000 × *g*, 4°C, 15 min). The cell pellet was resuspended in 20 mL of buffer F (50 mM sodium phosphate, 200 mM NaCl, 10% (v/v) glycerol, pH 8.0) and the cells were ruptured by sonication (3 × 10 sec, Soniprep 150, maximum power output). The cell lysate was clarified by centrifugation (11,000 × *g*, 4°C, 30 min), and the supernatant was filtered through a 0.45 μm filter. The filtered lysate was then loaded onto an Ni-NTA column (2 mL packed resin bed volume) and the column was washed overnight at 4°C with buffer F containing 10 mM imidazole. The His_6_-tagged proteins were eluted with buffer F containing 250 mM imidazole. The purified protein was mixed with His_6_-tagged TEV-protease and dialyzed overnight at 4°C against 2 L buffer G (20 mM Tris-HCl, 50 mM NaCl, 5% (v/v) glycerol, pH 7.5). The proteins thus released were cleaned up by batch extraction in a slurry of Ni-NTA resin equilibrated in buffer G. AceK_EC_ was expressed and purified as above, but from a modified pET-15b vector (a generous gift from Prof. Jia)^29^.

### Gel filtration and AUC

#### GFC

Analytical gel filtration with multi-angle light scattering was carried out using a 300 × 7.8 mm TSK-Gel G3000 SWXL column (Toso Haas) equilibrated with 150 mM NaCl, 20 mM Tris-HCl (pH 8.0) operating at a flow rate of 0.5 mL min^-1^. The column eluate was monitored in-line with a Mini-DAWN light scattering detector (λ = 690 nm), a quasi-elastic light scattering detector, differential refractometry and an absorption detector (280 nm). Molecular masses were calculated using Astra Software (Wyatt Technologies) and the Debye fit method^30^. A gel filtration marker kit (29 -700 kDa range) from Sigma-Aldrich was used to confirm accuracy of the measured masses.

#### AUC

Protein samples were first dialysed against 100 mM NaCl, 25 mM Tris-HCl (pH 7.5) to remove glycerol and DTT. AUC was performed using a Beckman Optima XL-I (AN-60 Ti rotor) fitted with absorbance and interference optics. Epon double-sector centrepieces were filled with 400 μL of sample solution or blank buffer and sedimented at 29160 × *g* for 24 h at 20°C. Absorbance data were acquired at λ = 280nm with time intervals of 2 min; interference scans were taken with time intervals of 1 min. Buffer viscosity, protein partial specific volumes and frictional ratios were calculated using SEDNTERP^31^. Data were analysed using SEDFIT^32^.

### Enzyme kinetic measurements

Kinetic analyses of ICL were carried out using two types of assay. In the “direct assay”, the enzyme was incubated in 25 mM imidazole, 5 mM MgCl_2_, 1 mM EDTA, 4 mM phenylhydrazine (pH 6.8) with a range of substrate concentrations (0 to 600 μM). The resulting glyoxylate-phenylhydrazone (ε_calc_ = 16,800 M^-1^ cm^-1^) was measured spectrophotometrically at λ = 324 nm. In the “coupled assay”, the enzyme was incubated in 50 mM MOPS-NaOH, 15 mM MgCl_2_, 5 mM DTT, 1 mM EDTA (pH 7.3) containing 400 μM NAD^+^ and 60 U of lactate dehydrogenase (LDH) from pig heart. The reduction of NAD^+^ to NADH (ε_calc_ = 6220 M^-1^cm^-1^) was measured spectrophotometrically at 340nm. All assays were carried out at 37°C.

The enzyme activity of IDH and ICD were measured using the same assay. The enzyme was incubated in 50 mM Tris-HCl, 5 mM MgCl_2_ (pH 7.5). To measure the K_M_ value for *iso*citrate, initial rates were measured across a range of (+)-potassium D-*threo*-*iso*citrate concentrations (0 to 600 μM) with a fixed concentration of NADP^+^ (200 μM). To measure the K_M_ value for NADP^+^, initial rates were measured across a range of NADP^+^ concentrations (0 to 600 μM) at a fixed concentration of *iso*citrate (200 μM). NADPH formation was recorded spectrophotometrically at λ= 340 nm (ε_calc_ = 6220 M^-1^cm^-1^). All assays were carried out at 37°C.

AceK_EC_ and AceK_PA_ kinase/phosphatase activity was assayed by coupling AceK activity to ICD activity as described previously^33^ with the following modifications. In a 96-well plate, 180 μL of phosphorylation reaction mixture (100 mM Tris-HCl, 1 mM ATP, 2 mM MgCl_2_, (pH 7.0) containing 10 μg ICD and 5 μg AceK) was incubated in each well for 1 h at 37°C. Following this, putative allosteric regulators were added to each reaction mixture to 5 mM final concentration. Each assay was carried out in triplicate. The reactions were then re-incubated for a further 30 min at 37°C to allow dephosphorylation of ICD and reactivation of ICD kinase activity. Aliquots (20 μL) of the reaction mixture were then transferred to a new UV-clear 96-well plate and 180 μL of reduction solution consisting of 100 mM Tris-HCl, 1 mM *threo*-D,L-*iso*citrate, 0.5 mM NADP^+^ and 2 mM MnCl_2_ (pH 7.0) was added to each well. The activity of ICD was detected spectrophotometrically by monitoring the rate of reduction of NADP^+^ at 340 nm using a FLUOstar Omega plate reader (BMG Labtech). As a control, ICD alone was assayed for its ability to reduce NADP^+^ in the presence of the putative allosteric regulators using the same method. Because phosphorylation of ICD by AceK inhibits the activity of ICD, higher ICD activity corresponds to increased AceK phosphatase activity. Confirmation of AceK_PA/EC_ phosphotransfer to ICD_PA/EC_ was also demonstrated using a [^32^P]-based phosphotransfer assay. The reaction conditions were identical to those described above, however, γ[^32^P]ATP (PerkinElmer) was used in place of unlabelled ATP. The proteins were resolved by SDS-PAGE and radiolabelling was revealed using a phosphor screen (GE Healthcare). Radioactivity was determined with a Typhoon PhosphorImager and quantitated with ImageQuant.

### Crystallisation conditions

Crystallisation conditions were screened then optimised using sitting drop vapor diffusion with 18, 17.5 and 10 mg/mL of purified protein for ICL, IDH and ICD, respectively. Crystals were obtained with a 1:1 protein : mother liquor mixture using a dragonfly^®^ discovery (ttp labtech) for mother liquor preparation and a crystallisation robot (mosquito^®^ HTS, ttp labtech) for liquid handling. ICL crystallised with 100 mM HEPES, 100 mM CaCl_2_, 20% (w/v) PEG 6000, 5% (v/v) glycerol, 1 mM glyoxylate (pH 5.0) and 2% thymol as additive. IDH crystallised in presence of 200 mM NaH_2_PO_4_, 21.5% (w/v) PEG 3350, 5% (v/v) glycerol and 150 μM NADP^+^. ICD crystallised in 100 mM sodium acetate, 30% (v/v) PEG 300 (pH 4.6). All crystals were grown for 2-6 days at 19°C then treated with cryoprotectant (24% (v/v) ethylene glycol and 76% (v/v) mother solution) before mounting in loops for data collection.

### X-ray Diffraction, Structure Determination, and Refinement

Diffraction data were collected remotely on the I02 or I04-1 beamline (as indicated in **Table S1**) at the Diamond Light Source Synchrotron (Didcot, UK). Diffraction data were processed using FastDP^27^, and the structures were determined by molecular replacement using Phaser^34^. ICL, IDH and ICD were solved using the structural templates 3I4E (to be published), 4ZDA (to be published) and 1BL5^35^ from *Burkholderia pseudomallei*, *Mycobacterium smegmatis* and *Escherichia coli*, respectively. Automated refinement was performed using Refmac5^36^. Manual modelling and refinement were performed in COOT^37^. Data collection and refinement statistics are listed in **Table S1.**

### Western blotting

Polyclonal antibodies were raised in rabbits against each of the purified proteins (Biogenes.De). The antisera were pre-absorbed against an acetone extract of a mutant strain defective in the protein of interest (e.g., the anti-ICD antisera were pre-absorbed against an acetone extract of a confirmed *icd* mutant). The cleaned-up antisera, appropriately diluted, were then used directly in Western assays. Western blots were developed using goat anti-rabbit secondary antibodies.

## Supplementary Information Figure Legends

**Figure S1.**
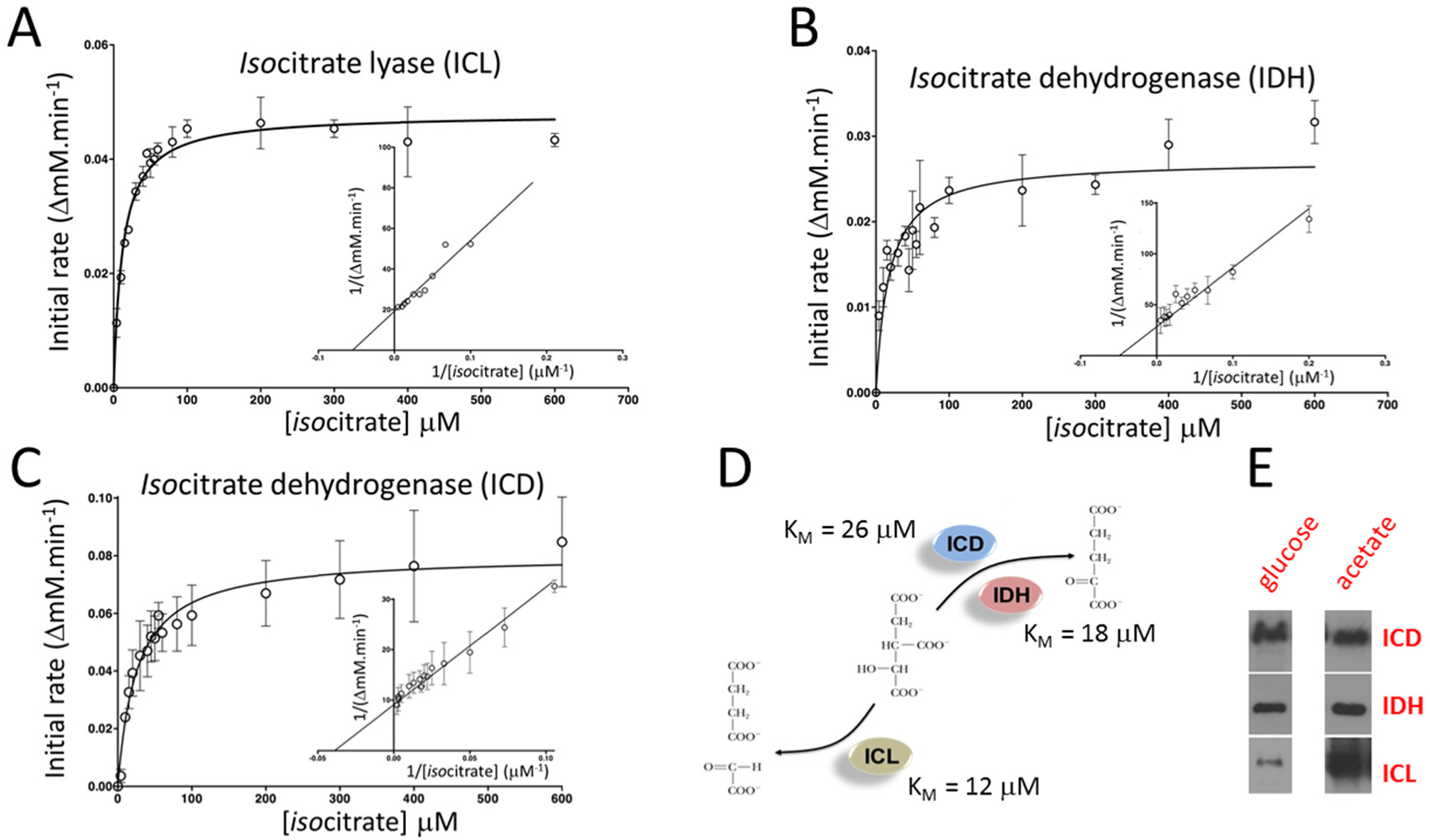
Enzyme kinetics (*iso*citrate dependency) of *P. aeruginosa* ICL **(A)**, IDH **(B)** and ICD **(C)**. The main body of each panel shows the direct plot and the insert shows the corresponding Lineweaver-Burk transformation. Extracted kinetic constants are shown in **Table I**. Each assay was performed in triplicate. Error bars correspond to ± 1 standard deviation. **(D)** Schematic showing the calculated K_M_ (*iso*citrate) of each enzyme and the reaction catalysed. **(E)** Semi-quantitative Western blot showing that ICD and IDH are expressed at similar levels during growth on glucose or acetate, and that ICL is expressed during growth on glucose but that the level of enzyme expression increases during growth on acetate.

**Figure S2.**
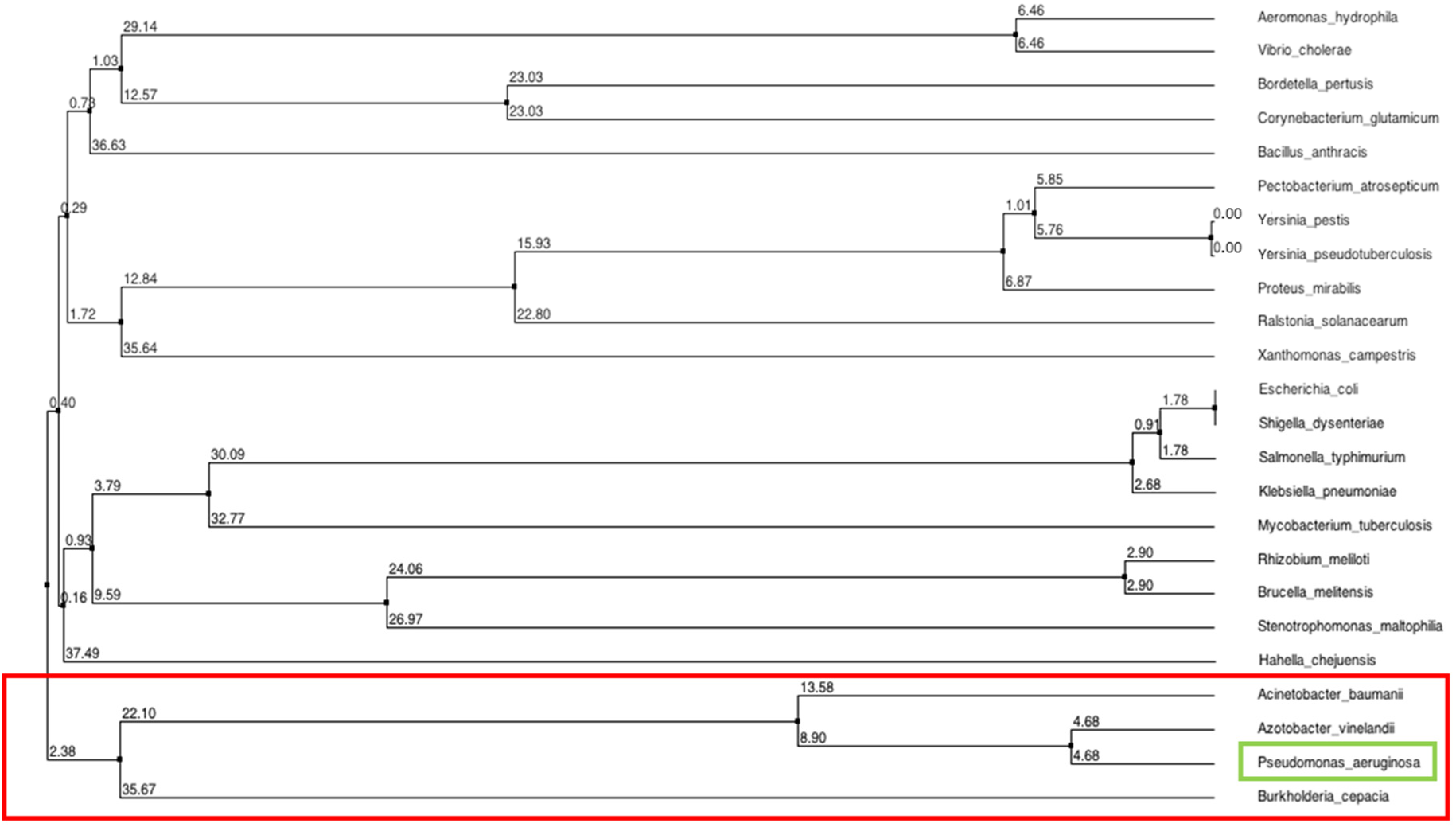
Phylogenetic tree of ICL from a subset of pathogens. Note how the *P. aeruginosa* ICL forms a distinct cluster (boxed in red) with ICL from *Acinetobacter baumanii*, *Azotobacter vinelandii* and *Burkholderia cepacia*. The tree was generated after alignment of all amino acid sequences using ClustalOmega and neighbour joining was calculated from the percentage sequence identity in JalView.

**Figure S3.**
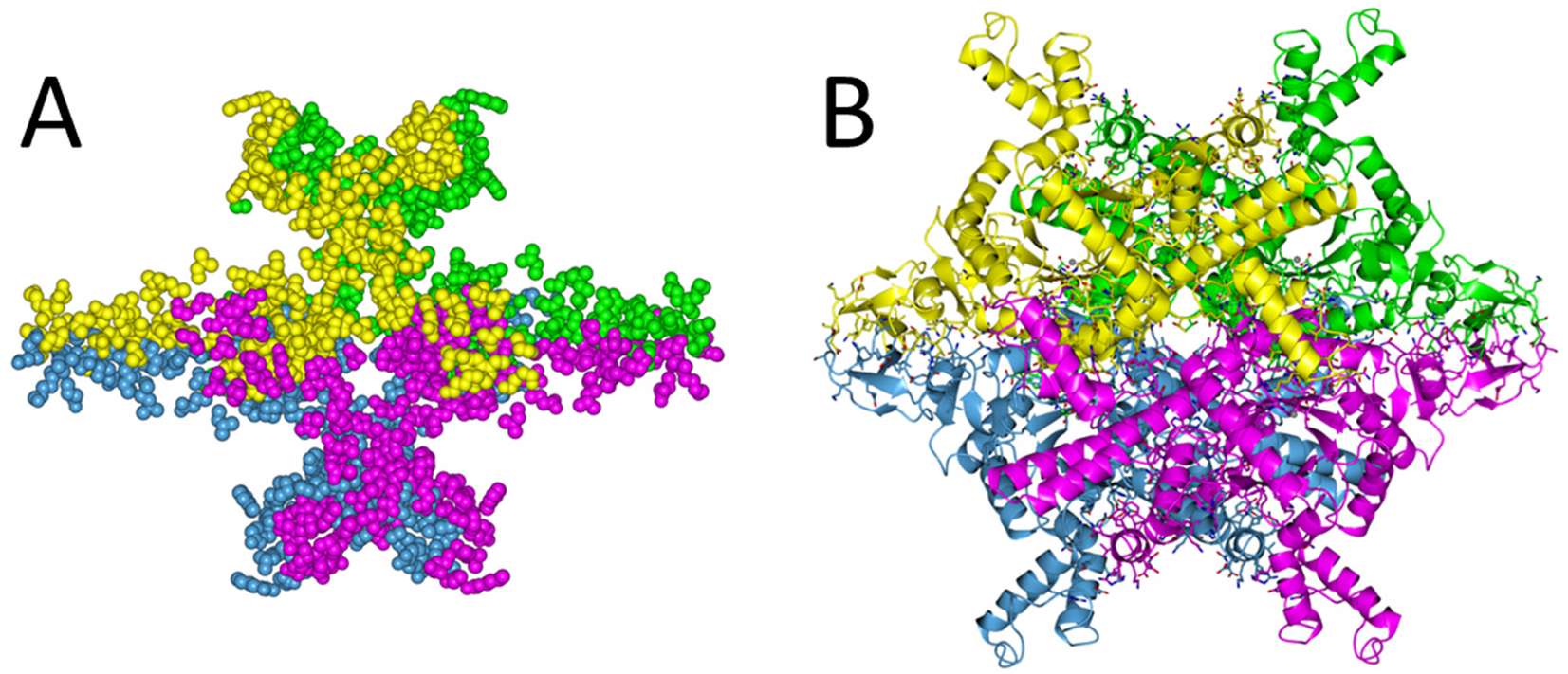
The *P. aeruginosa* ICL tetramer is held together by numerous inter-subunit bonds contributed by each protomer. **(A)** Space-filling view of the residues involved in tetramerisation of *P. aeruginosa* ICL. **(B)** The same residues represented as sticks supported by each protomer of the tetramer. The data shown are based on a COCOMAPS estimation, and the total excluded surface area is 12,352 Å^2^.

**Figure S4.**
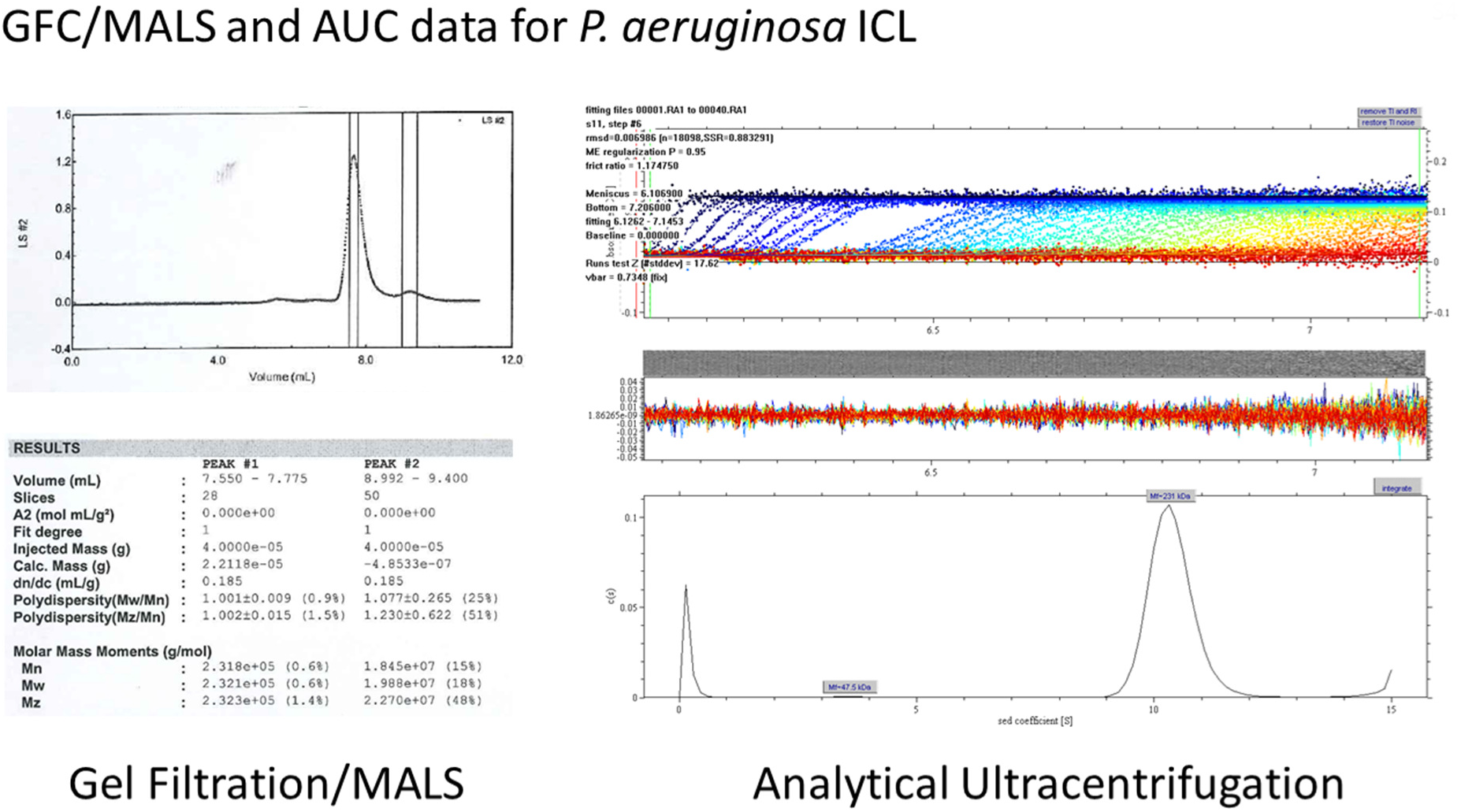
Solution structure of ICL determined by gel filtration chromatography coupled with multi-angle light scattering (GFC-MALS) and analytical ultracentrifugation (AUC). The overall molecular mass of ICL inferred from GFC-MALS analysis was 232 kDa. AUC analysis yielded a very similar estimated molecular mass of 231 kDa. Given that the calculated molecular mass of one ICL polypeptide is 59 kDa, *iso*citrate lyase is likely to be a tetramer (M_w_ = 236 kDa) in solution.

**Figure S5.**
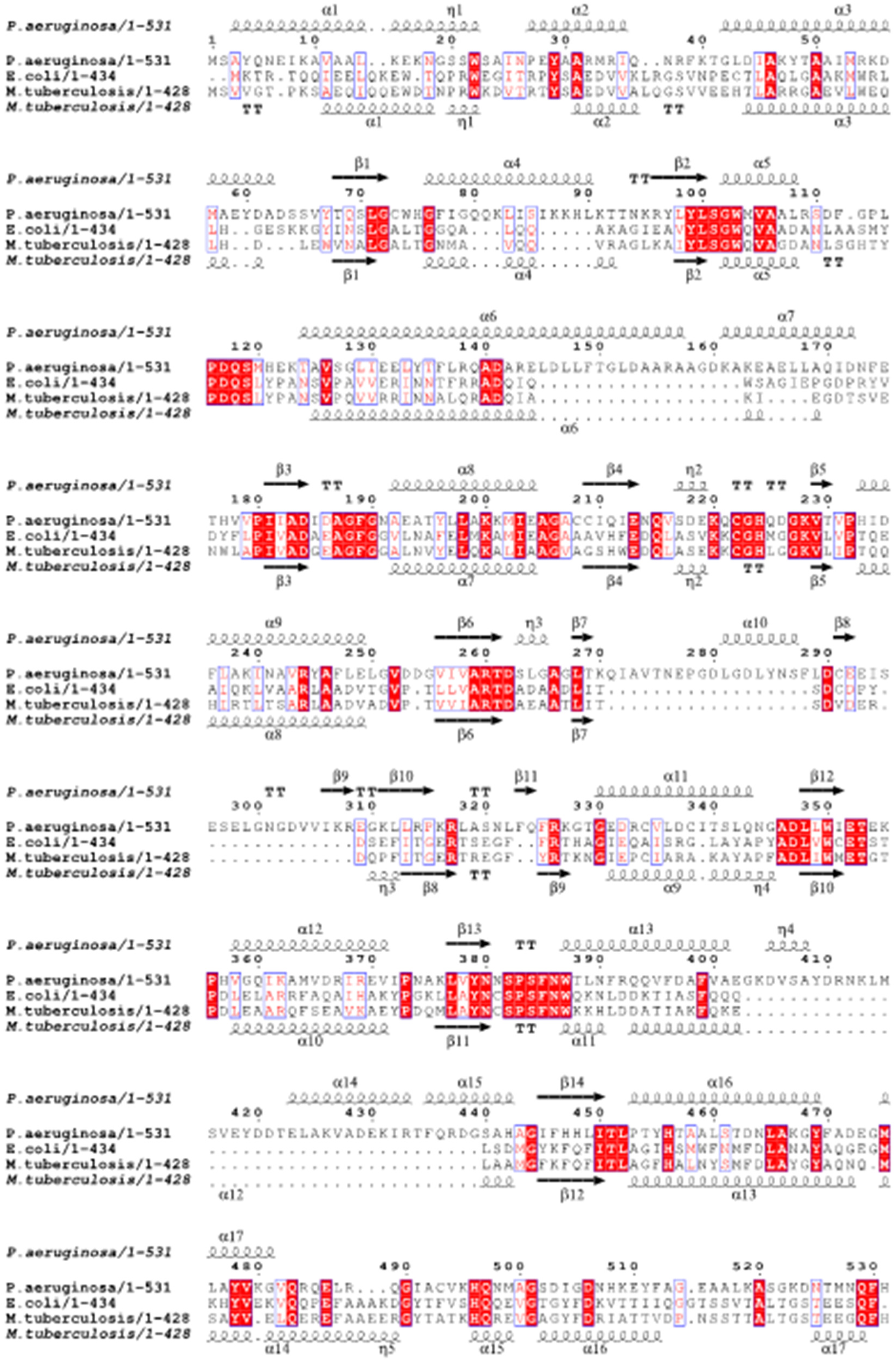
ESPript alignment and secondary structure analysis of *iso*citrate lyase from *P. aeruginosa*. The amino acid sequences of ICL from *P. aeruginosa*, *E. coli* and *M. tuberculosis* are shown, along with the corresponding secondary structure assignments (*P. aeruginosa* on the top row and *M. tuberculosis* (PDB 1F8I) on the bottom row).

**Figure S6.**
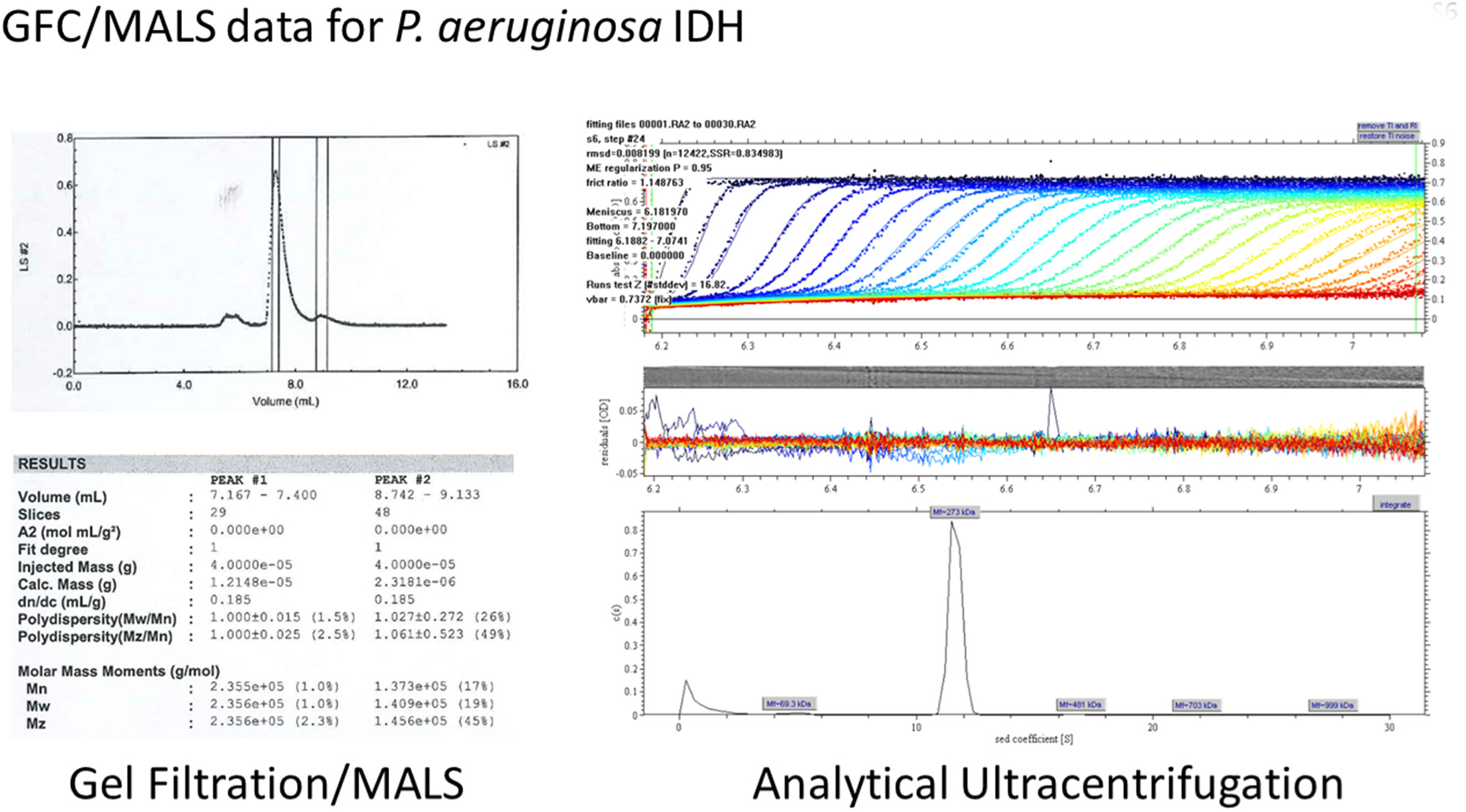
Solution structure of IDH determined by gel filtration chromatography coupled with multi-angle light scattering (GFC-MALS) and analytical ultracentrifugation (AUC). The overall molecular mass of IDH inferred from GFC-MALS analysis was 236 kDa. AUC analysis yielded a higher estimated molecular mass of 273 kDa. Given that the calculated molecular mass of one IDH polypeptide is 82 kDa, these data suggest that IDH could be a trimer or elongated dimer in solution. However, we also note that the protein has a rather low frictional coefficient (*f* = 1.15), consistent with it adopting a compact, globular configuration.

**Figure S7.**
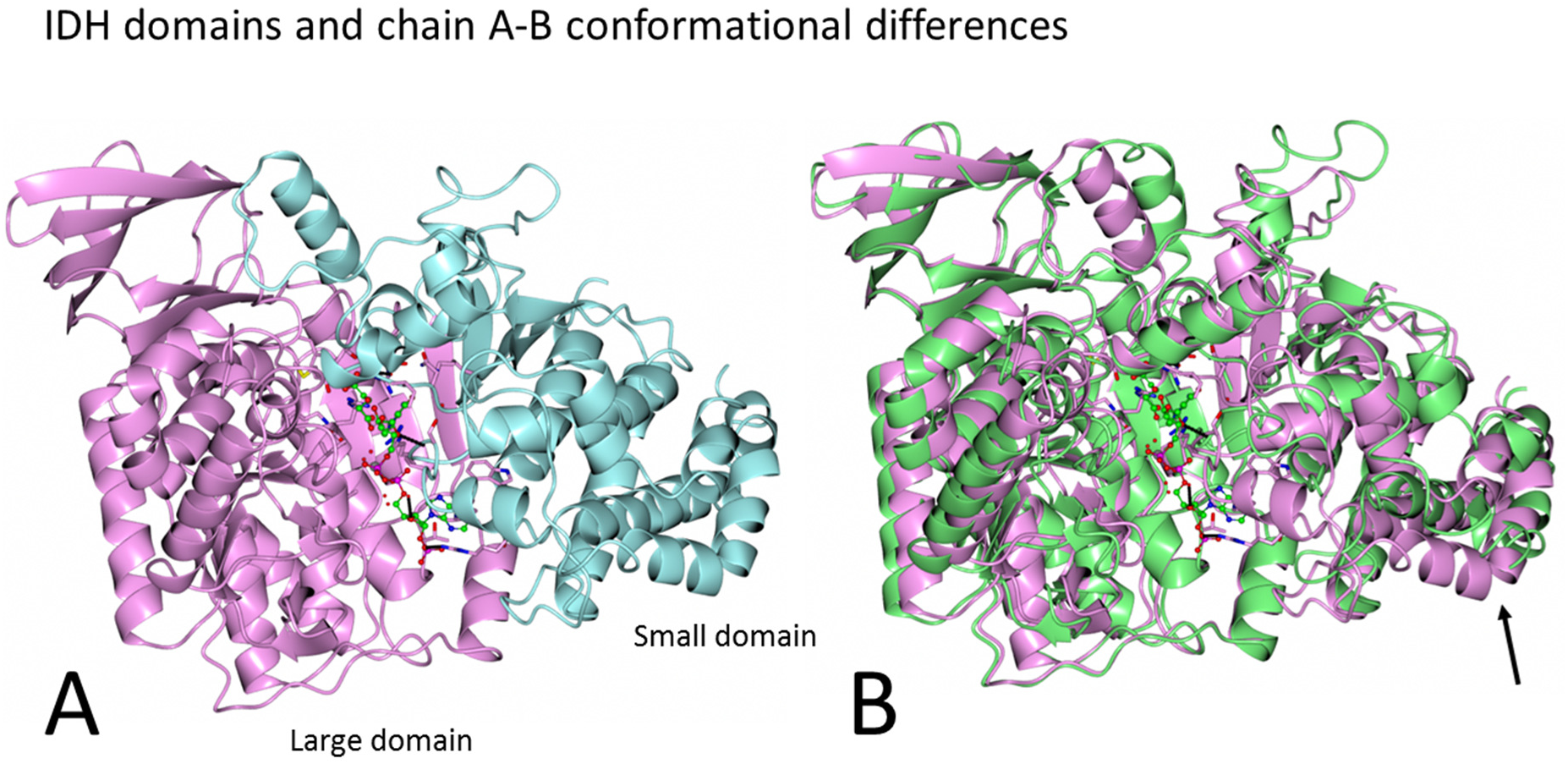
IDH domains and conformational differences in the presence of bound reactions products. The figure shows a cartoon representation of *P. aeruginosa* IDH. **(A)** IDH is comprised of two domains; a large one (mauve) spanning Leu139 - Leu571, and a small one (teal) spanning Ser5 - Val138 and Met572 - Ala741. **(B)** Superposition of IDH chain A (mauve) and chain B (light green). The conformational differences between each polypeptide chain affect most parts of the molecule, although they are particularly obvious in the small domain in the region marked by the arrow.

**Figure S8.**
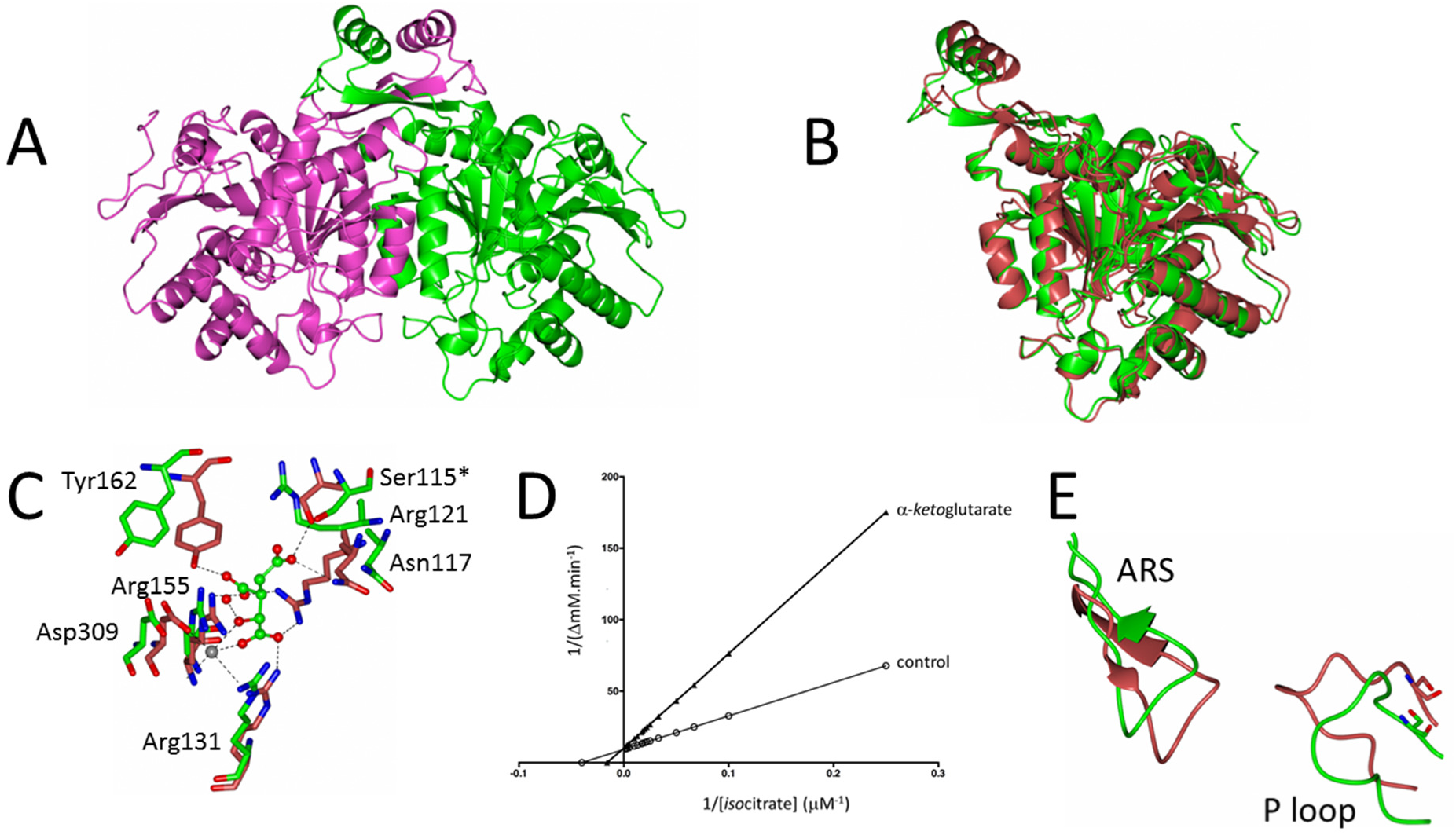
Structure and kinetics of *P. aeruginosa iso*citrate dehydrogenase (ICD). **(A)** Ribbon representation of the ICD dimer. The two protomers are highlighted in light purple and green. Dimerization is largely mediated by the clasp domain, shown here at the top of the structure, although additional contacts between helices at the core of the structure also play a role. **(B)** The overall structure of *P. aeruginosa* ICD is very similar to that of *E. coli* ICD. The panel shows a superposition of the *P. aeruginosa* structure (green, PDB 5M2E) and the *E. coli* ICD structure (red, PDB 4AJA). The only significant structural difference between the proteins is that in the projection forming the clasp-like dimerization determinant, the *P. aeruginosa* structure forms a large β-strand (encompassing residues Cys196 to Ser204) that is absent in the *E. coli* protein. Overall, the secondary structures superimpose with an RSMD of 1.94 Å. **(C)** The known catalytic residues in *E. coli* ICD_EC_ (red, PDB 4AJA) are conserved in *P. aeruginosa* ICD (green, PDB 5M2E). The ball-and-stick molecule of *iso*citrate and the Mg^2+^ (gray) shown in this panel are modelled from the *E. coli* ICD structure, since these are not present in the *P. aeruginosa* ICD structure. The absence of substrate in the *P. aeruginosa* apo structure likely explains why not all of the side chains point inwards to coordinate the *iso*citrate. **(D)** ICD is feedback inhibited by α-*keto*glutarate. A Lineweaver-Burk analysis indicates that the inhibition is competitive. **(E)** The AceK recognition motifs are conserved in the *P. aeruginosa* ICD structure. The configuration of the ARS and P-loop are similar in ICD_PA_ (green) and ICD_EC_ (red).

**Figure S9.**
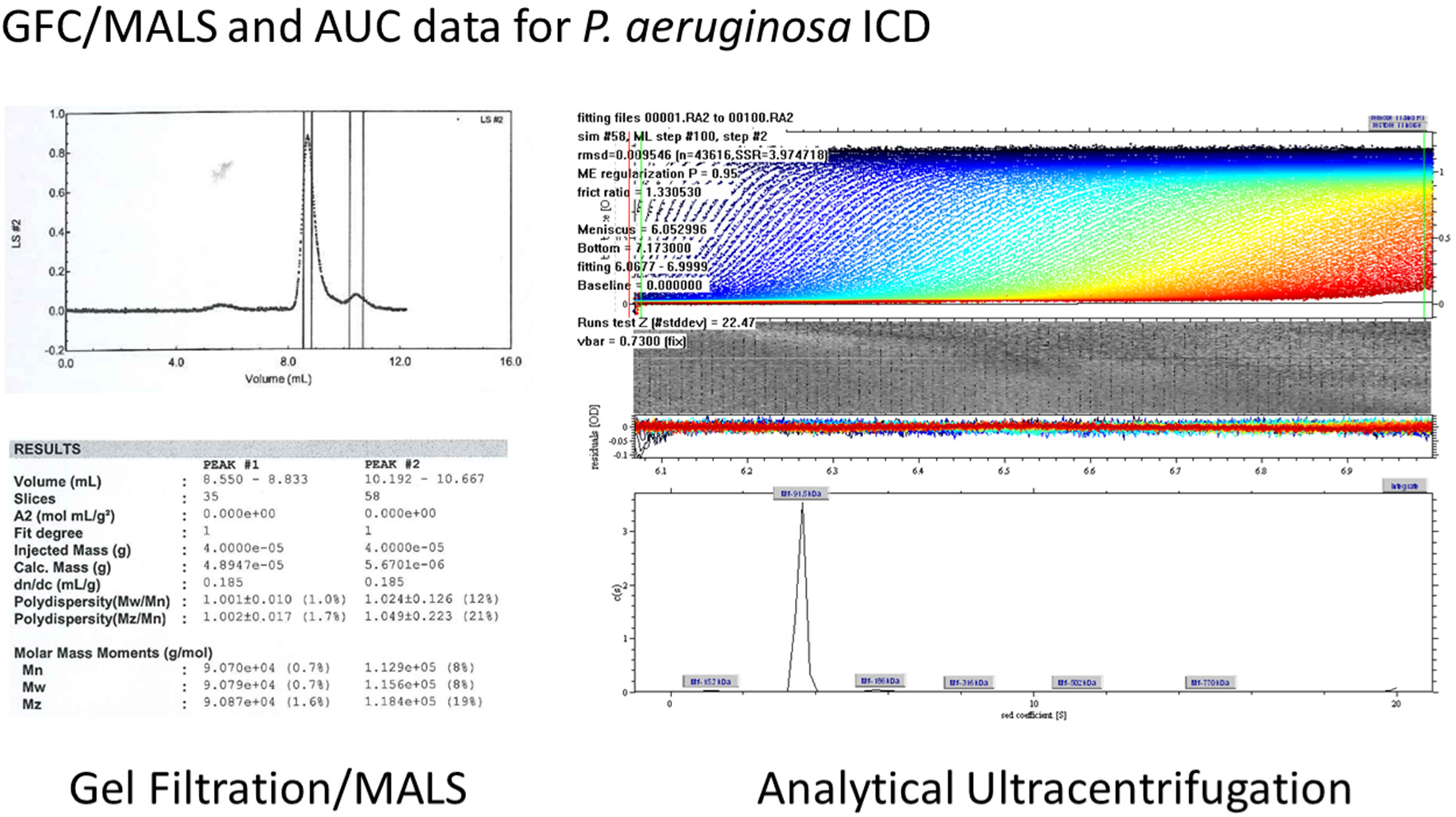
Solution structure of ICD determined by gel filtration chromatography coupled with multi-angle light scattering (GFC-MALS) and analytical ultracentrifugation (AUC). The overall molecular mass of ICD inferred from GFC-MALS analysis was 90 kDa. AUC analysis yielded a very similar estimated molecular mass of 91.5 kDa. Given that the calculated molecular mass of one ICD polypeptide is 45 kDa, the ICD *iso*citrate dehydrogenase is likely to be a dimer (M_w_ = 90 kDa) in solution.

**Figure S10.**
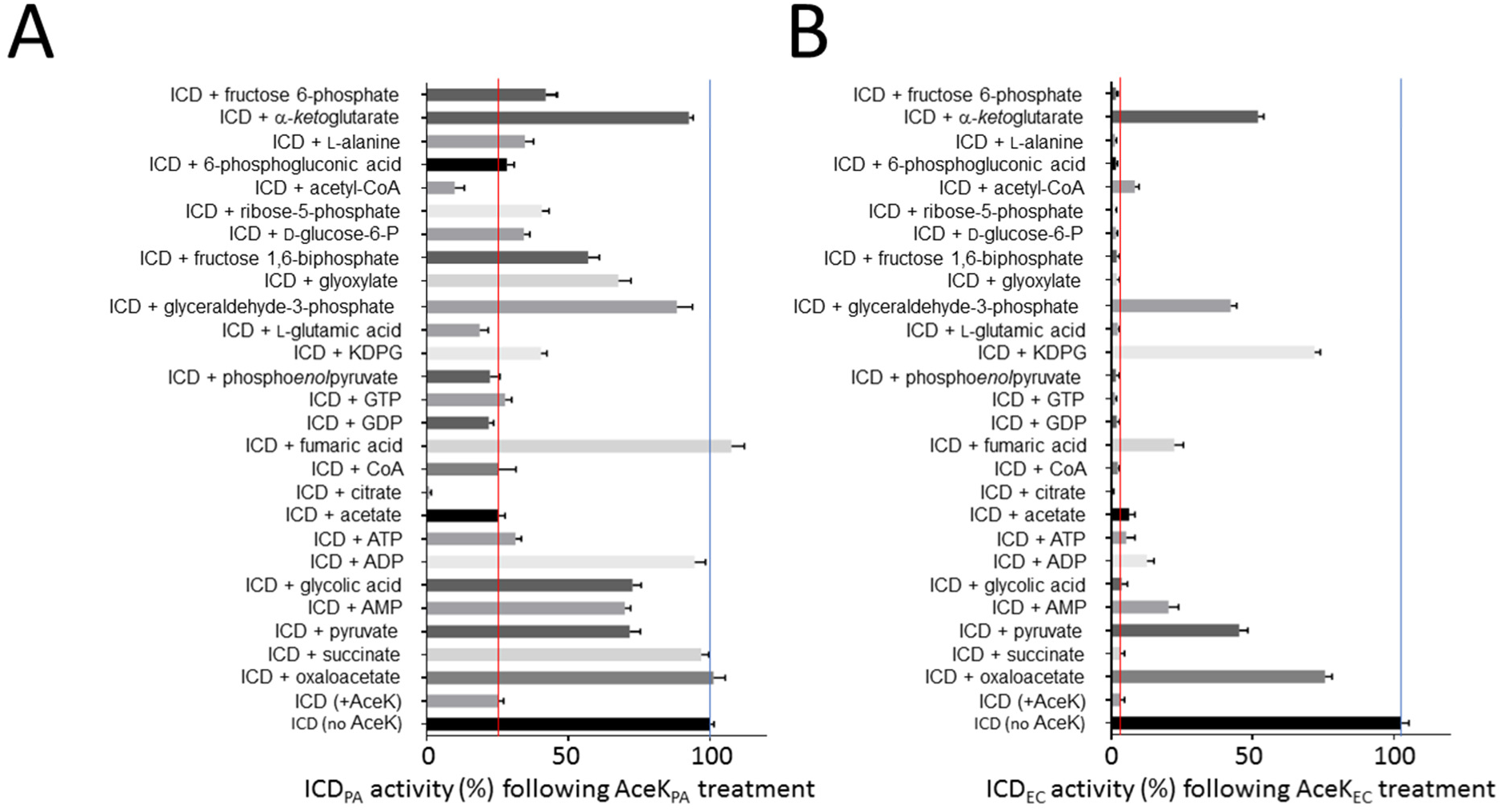
Impact of small molecule regulators of AceK kinase/phosphatase activity. **(A)** ICD_PA_ was first inactivated by incubation for 60 min with AceK_PA_ and ATP. Following this, the indicated small molecule regulators were added (5 mM final concentration in all cases) and the reaction was allowed to continue for a further 30 min. The *iso*citrate dehydrogenase activity of each sample was then measured. Note how ICD retains about 25% of its activity even after extensive treatment with AceK, unless acetyl-CoA or citrate are present (see main text for details). The blue line represents the activity of ICD that has not been treated with AceK and the red line indicates the activity of ICD that has been treated with AceK (but not further treated with potential small molecule regulators). **(B)** A similar experiment to that just described was also carried out with ICD_EC_ with AceK_EC_. The *E. coli* AceK more profoundly inhibits *E. coli* ICD activity compared with the *P. aeruginosa* homologue AceK/ICD pair. All plots were generated using GraphPad Prism 6, and the errors bars correspond to ± 1 standard deviation. All experiments were performed in triplicate.

**Figure S11.**
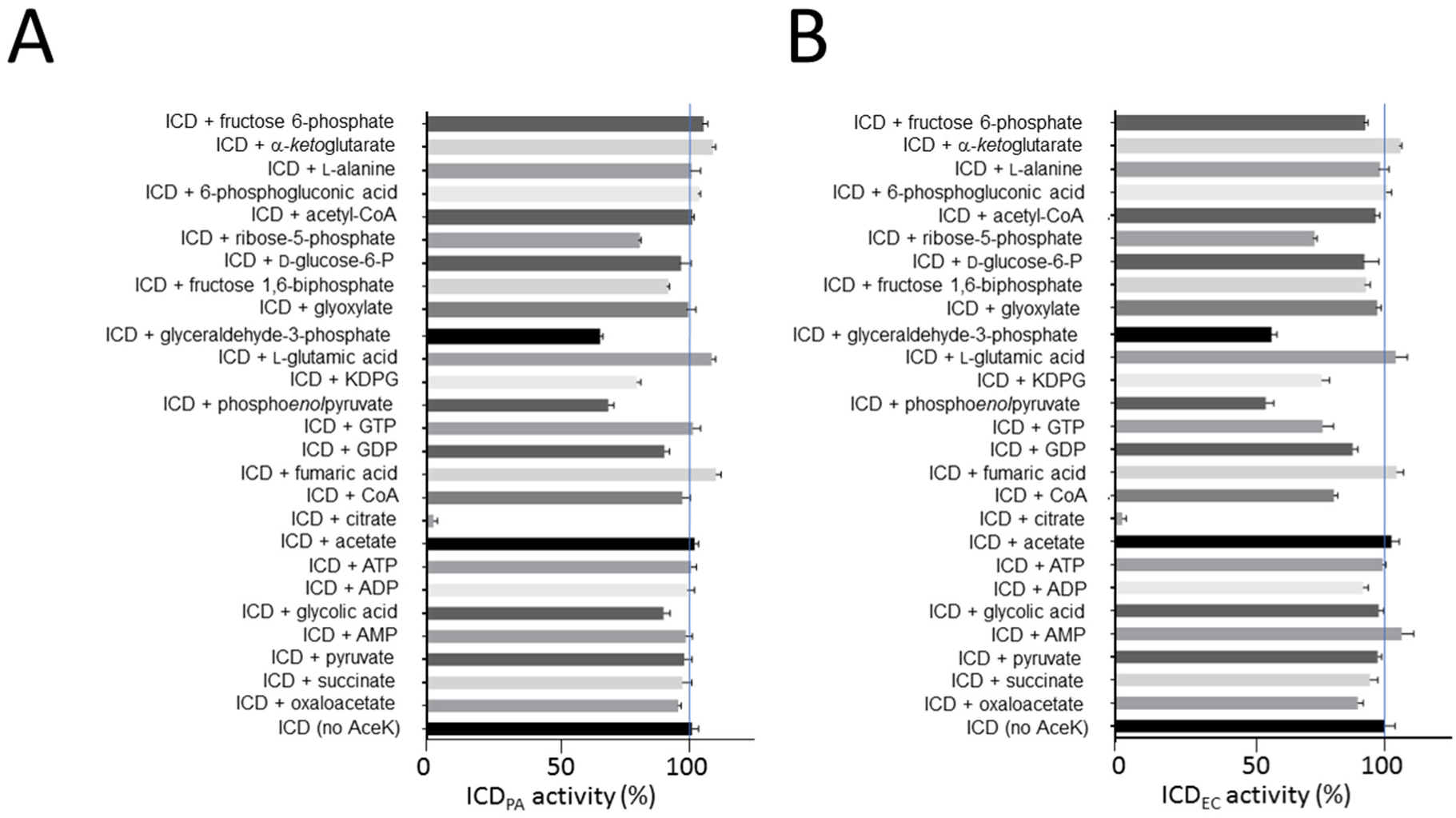
ICD *iso*citrate dehydrogenase activity is not intrinsically affected by most of the regulators that impact on AceK. **(A)** The metabolites tested as putative AceK regulators in **Figure S10** were also evaluated for their ability to directly activate/inhibit *P. aeruginosa* ICD. Note that ICD retains almost full activity in the presence of most of these compounds, although PEP, glyceraldehyde 3-phosphate, 2-*keto*-3-deoxyphosphogluconate, and ribose 5-phosphate slightly depress ICD activity when present at 5 mM, and citrate blocks ICD activity completely. **(B)** The same experiment as in (A) carried out with ICD_EC_.

**Table S1.** Crystallographic statistics.

|  | ICL | IDH | ICD |
| --- | --- | --- | --- |
| **Data collection statistics** |  |  |  |
| Radiation source | Diamond (UK), I02 | Diamond (UK), I04-1 | Diamond (UK), I04-1 |
| Wavelength (Å) | 0.9795 | 0.9282 | 0.9282 |
| Space group | I222 | C222<sub>1</sub> | P12<sub>1</sub>1 |
| Cell dimensions |  |  |  |
| a, b, c (Å) | 80.94, 116.02, 128.53 | 126.46, 149.02, 201.13 | 88.88, 95.55, 104.01 |
| α, β, γ (°) | 90, 90, 90 | 90, 90, 90 | 90, 99, 90 |
| Resolution range (Å) | 1.88-29.49 (1.88-1.93) | 2.71-29.62 (2.71-2.78) | 2.70-47.78 (2.70-2.79) |
| Total reflections | 614357 (27689) | 940727 (66819) | 304691 (17419) |
| Unique reflections | 48986 (3287) | 51799 (3682) | 44233 (2978) |
| Multiplicity | 12.5 (8.4) | 18.2 (18.1) | 6.9 (5.8) |
| Completeness (%) | 99.4 (92.1) | 99.6 (97.2) | 98.0 (89.4) |
| Mean I/sigma(I) | 21.7 (3.0) | 17.3 (3.7) | 17.9 (2.0) |
| R-merge | 0.086 (0.656) | 0.176 (0.891) | 0.068 (0.826) |
| **R-pim** | 0.036 (0.340) | 0.059 (0.304) | 0.028 (0.348) |
| CC-half | 0.99 (0.78) | 0.99 (0.92) | 0.99 (0.79) |
| **Refinement** |  |  |  |
| R<sub>work</sub> (%) | 18.32 | 20.80 | 24.56 |
| R<sub>free</sub> (%) | 21.17 | 26.67 | 28.37 |
| **No. of molecules per ASU** | 1 | 2 | 4 |
| No. of total atoms (no H) | 3798 | 11551 | 12832 |
| atoms for ligands | 6 | 58 | n/a |
| atoms for waters | 182 | 197 | 36 |
| **Ramachandran plot analysis, number of residues in;** |  |  |  |
| Favoured regions (%) | 96.90 | 93.91 | 92.19 |
| Allowed regions (%) | 2.69 | 5.54 | 7.03 |
| Disallowed regions (%) | 0.41 | 0.55 | 0.78 |
| **B-factor (Å<sup>2</sup>)** |  |  |  |
| Average/Wilson | 35.10/26.42 | 52.31/43.36 | 114.21/68.00 |
| Ligands | 52.86 | 28.88 | n/a |
| Solvent | 32.55 | 35.87 | 38.04 |
| **RMS deviations** |  |  |  |
| Bond lengths (Å) | 0.01 | 0.01 | 0.003 |
| Bond angles (°) | 1.43 | 1.52 | 0.602 |
| **PDB entry** | **6G1O** | **6G3U** | **5M2E** |

